# Revisiting the evidence for long-lived balancing selection in humans

**DOI:** 10.1101/2025.11.10.687682

**Authors:** Hannah Munby, Molly Przeworski

## Abstract

Balancing selection maintains variation in a population and, when sufficiently long-lived, can lead to polymorphisms shared identical by descent across species. In settings where sharing is extremely unlikely to occur under neutrality, such trans-species polymorphisms provide clear-cut evidence for balancing selection. These considerations have motivated efforts to identify trans-species polymorphisms between humans and close evolutionary relatives, such as chimpanzees, and have led to the discovery of a handful of examples. However, recent demographic modeling suggests that gene flow between humans and chimpanzees may have persisted until much more recently than previously thought, implying that many trans-species polymorphism could potentially have arisen by chance. Here, we reassess the evidence for long-term balancing selection in humans by characterizing shared single base pair polymorphisms in whole genome sequences from 2504 humans and 59 chimpanzees. Using recently-developed methods to reconstruct local gene genealogies, we infer the evolutionary histories surrounding the oldest shared variants and distinguish true trans-species polymorphisms from recurrent mutations, recovering both previously known and novel examples. We then examine multiple lines of independent evidence for the action of natural selection on this set of trans-species polymorphisms. These analyses suggest that the vast majority may have arisen by chance, in regions of neutral, incomplete lineage-sorting. In other words, while a subset of such loci remain compelling candidates for long-term balancing selection, the finding of trans-species polymorphism shared between humans and chimpanzees is not, in itself, strong evidence for long-lived balancing selection.

## Introduction

Genetic differences among individuals are shaped by both neutral and selective forces. In studying these genetic differences, much attention has been given to the study of directional selection, which favors a single genetic allele over others and acts to reduce variation at linked sites [1,2]. But it is also possible that variation is in itself adaptive, such that multiple alleles at a single locus are maintained in the population over time by natural selection [3]. There is a long history of theoretical work describing various selective regimes that result in the maintenance of diversity at a locus longer than expected under genetic drift alone, collectively referred to as balancing selection, and their effects on linked neutral diversity [4,5]. Cases of balanced polymorphisms include the S-locus, which leads to self-incompatibility in certain families of flowering plants [6], the disease resistance R-locus in *Arabidopsis [7]*, structural polymorphisms that control sexual reproduction strategies in the walnut family [8,9], and balanced inversions that maintain behavioral ecomorphs in animals, such as the three male ruff morphs [10]. Examples are few, however, and the broader questions of how much variation is actively maintained by selection, the types of the variants that are maintained, why they have been selected and over what timescales they have persisted, all remain largely open [11].

In humans specifically, despite millions of genomes now sequenced, we have only relatively few well-characterized examples of balancing selection, most of which seem to be due to heterozygote advantage. Perhaps most famously, in geographic regions endemic for *P. falciparum* malaria, individuals with one sickle allele and one wild-type allele have a higher fitness than either homozygote [12]. In this case, the selection pressures are known: in the heterozygous state, the sickle cell allele confers resistance to malaria but in the homozygous state, the fitness benefit provided by protective effect of the allele is outweighed by it leading to sickle cell disease. The sickle cell allele is thus maintained by strong balancing selection in these geographic regions. This polymorphism is thought to have arisen very recently in human evolution, however, only around 7000 years ago [13]. Other examples of heterozygote advantage are again thought to be in response to malaria and are similarly young (e.g., α-thallassemia [14] and G6PD deficiency [15]).

The recency of these examples is perhaps not surprising. Heterozygote advantage is unlikely to be evolutionary stable, given that in each generation, there is a segregation load of heterozygous individuals having less fit homozygous offspring [16]. This load can ultimately be resolved by a gene duplication that fixes both alleles in the genome, as is hypothesized to have occurred for opsins for example [17,18]. Modeling suggests that, in contrast, other modes of balancing selection–including frequency-dependent selection, temporally or spatially-varying selection, and selection driven by interaction with other species–have the potential to maintain alleles for much longer [7,19,20]. While such long-lived balanced loci are expected to be rarer, they are the result of strong and persistent selective forces and are thus of importance in understanding adaptation, as well as in identifying genomic regions in which variation is of functional importance.

The identification of long-lived balanced polymorphisms often relies on the impact that selection has on the underlying genealogy, specifically on the fact that balancing selection leads the time to the most recent common ancestor (TMRCA) at the target locus to be older than expected under neutrality (or purifying selection). This property causes a signal of increased diversity at neutral sites linked to the selected locus, an observable pattern relied on by many approaches to detect selection in genomic data [21]. The exact distortion of the underlying genealogy can depend on the mode of balancing selection: as an example, the effects of fluctuating selection can be distinct from those of overdominance or frequency-dependent selection [22–24].

Methods tend to assume a constant size, panmictic population, even though changes in population size and population structure will also shape the genealogy [25–27]. More generally, existing approaches to detect balancing selection tend to rely on strong assumptions about demography and the mode of balancing selection, as well as about mutation and recombination rates. For example, likelihood-based methods (e.g., [28–30]) assume a null model of neutrality under a known population history and an alternative model of heterozygote advantage under a constant population size. Moreover, when such methods rely on a model of heterozygote advantage, they model the balanced polymorphism at deterministic equilibrium allele frequencies, which may not capture the dynamics under alternative forms of balancing selection [31,32]. Other model-based methods make fewer assumptions about the mode of selection but assume the data come from an unstructured population (e.g., [33]) and are sensitive to model misspecification [11]. To overcome these limitations, one approach is to focus on regions of the genome with the highest inferred TMRCAs, which are plausibly enriched for long-lived balanced polymorphisms [34]; however, the lack of a clear null distribution to compare against makes it difficult to draw firm conclusions about whether these tail cases are truly targets of selection.

There is, however, a special case thought to provide unequivocal evidence for balancing selection [35]: when a polymorphism persists long enough for a variant that arose in an ancestral species to be maintained to the present day, and thus leads to allele sharing between relatively divergent species through identity by descent (IBD). In humans, as well as in other species, such trans-species polymorphisms (TSPs) with chimpanzees or more distant evolutionary relatives have long been held as a gold standard [4,36], in large part because modeling suggested that it would be highly unlikely for alleles to be shared IBD between humans and other primate species by genetic drift alone [36]. Moreover, well-established examples of balancing selection, such as occurs at loci in the Major Histocompatibility Complex (*MHC*) [37] and at the *ABO* gene, show the clustering by allele rather than by species expected from IBD: for example, in *ABO*, a small genomic region differs more between humans carrying alleles A versus B than it does between humans and gibbons carrying allele A [35]. Beyond these two cases, only a handful of TSPs are known between humans and other primate species [38–40]. Nonetheless, evidence that a number of these examples are under selection due to host-pathogen interactions [40–42], as well as theory suggesting such interactions can maintain host variation [43], have led to the hypothesis that host-pathogen immune functions are the main force driving long-term balancing selection in great apes [44].

The interpretation of variation shared IBD between humans and chimpanzees as strong evidence of long-term balancing selection rests on the assumption that such a genealogical pattern should be exceedingly rare under neutrality: that a neutral polymorphism is very unlikely to survive without being lost or fixed across the tens of million years between the two species. Recent findings about human demographic history weaken this assumption. Reanalysing effective population size histories using high quality telomere-to-telomere hominid assemblies, Cousins et al. 2026 inferred deep coalescence times in a small, but non-trivial, fraction of both the human and chimpanzee genome, with roughly 1% of the genome remaining uncoalesced by 5 Mya in humans and 0.5% in Western chimpanzees [45]. Additionally, they inferred a coincident elevation in effective population size around 3-6 Mya in humans, chimpanzees, and bonobos (but not orangutan or gorilla). From these results, they concluded that the separation of the *Homo* and *Pan* lineages was not a clean split from a panmictic ancestor but a protracted, "complex" speciation event, with a structured ancestral population and descendant groups that continued to exchange migrants after their initial divergence. If a substantial portion of the genome remains uncoalesced until the ancestral population in both species, neutral incomplete lineage sorting may be far more common than previously appreciated. Viewed in another way, a more recent final split time between humans and chimpanzees implies a shorter time period over which variants must be maintained until the present, making this configuration more likely by chance. Thus, these results raise the possibility that at least some trans-species polymorphisms (TSPs) shared between humans and chimpanzees may have persisted without balancing selection.

In light of these findings, we sought to reassess the evidence for ancient balancing selection by systematically identifying trans-species polymorphisms between humans and chimpanzees, then evaluating the evidence for their maintenance by natural selection. Since the last comprehensive inventory of such variants over 10 years ago based on 10 chimpanzee sequences at medium coverage, there have been many methodological and empirical developments. In particular, we use whole genome sequences from 2504 humans [46] and 59 chimpanzees sequenced at ∼30X, including 10 chimpanzee genomes that we resequenced, to identify the common variants shared between the two species (see Methods).

This set will include trans-species polymorphisms but will largely consist of recurrent mutations. Reliably distinguishing between the two requires an accurate reconstruction of the local gene genealogy. A challenge in identifying TSPs in humans is that the footprint around such an old site is expected to be on the order of 100-1000 base pairs (bps) for human recombination rates [36], meaning that examples can be missed if tree reconstructions are based on larger window sizes [36]. In recent years, however, scalable ancestral recombination graph (ARG) reconstruction methods have been developed that allow for the inference of all local genealogies along the genome [34,47–49]. These methods also infer mutational ages, and have been used to do so in large samples of humans [47,48]. We use these allele ages to narrow our focus to variants that are plausibly old enough to be shared IBD with chimpanzees. By performing ARG reconstruction on a combined sample of the two species, we identify the subset that are likely to be TSPs. With this set in hand, we consider population genetic and functional evidence about them in order to assess whether, as a set, the TSPs are more likely to have resulted from balancing selection or neutral processes.

## Results

### Characterizing common variation shared between humans and chimpanzees

In order to identify trans-species polymorphisms, our first step is to characterize the set of common variants shared between humans and chimpanzees. To survey genetic variation in chimpanzees, we compile 49 publicly available whole-genome sequences from population sequencing [50] and three pedigree studies [51–53]. We also include 10 newly sequenced samples from a three-generation pedigree that we had collected for other purposes, leading to a total of 59 individuals. We remove first- and second-degree relatives, leaving 43 samples assigned to one of three chimpanzee subspecies on the basis of prior information and genetic similarity: 15 western (*P. t. verus*), 14 central (*P. t. troglodytes*) and 14 eastern (*P. t. schweinfurthii*) (see Methods and Supplementary Figure 1).

We jointly call and filter genotypes for these samples using the latest chimpanzee reference genome (PanTro6) before lifting variants to the human reference. Specifically, we reciprocally lift variants (and the surrounding ±100 bp) between the two assemblies, keeping only those that map consistently. Because we are interested in long-term balancing selection, and over long time periods rare variants tend to be lost by drift, we focus on common variants: SNPs with a minor allele frequency of at least 5% in the 2,504 unrelated humans of the 1000 Genomes Project phase 3 (1000GP) and a minor allele count of three or more in our 43 unrelated chimpanzees. We further restrict our analysis to the autosomes. Intersecting these two sets and removing variants in regions with known copy-number variants and segmental duplications (see Methods), we identify 66,650 biallelic autosomal SNPs that are polymorphic and present at appreciable frequency in both species.

We annotate variants with allele-age estimates previously obtained by applying two local genealogical reconstruction methods, *Relate* [47] and *SINGER* [48], to 1000GP data. *Relate* has been shown to underestimate the age of variants that lie in regions with particularly ancient genealogies [54]. While *SINGER* appears substantially less prone to this bias [48], there exist age estimates for far fewer of our shared SNPs. We therefore rely on *Relate* as our primary source, having confirmed that it yields similar results to *SINGER* for SNPs for which both are available. *Relate* estimates the ages for each human population. We use those for the Yoruba in Ibadan (YRI), because African populations have deeper coalescent histories, which should help to better resolve the ages of old alleles; YRI is also among the largest 1000GP samples, and using it aids comparison with previous work [38]. We note that genealogical reconstruction methods do not provide estimates of mutations, but instead infer the start and end times of the branch on which each mutation is inferred to have arisen. Assuming neutrality, the mutation age is uniformly distributed along that branch [55]. We use this assumption to compute the expected number of SNPs older than a given age (Figure 1A).

**Figure 1:**
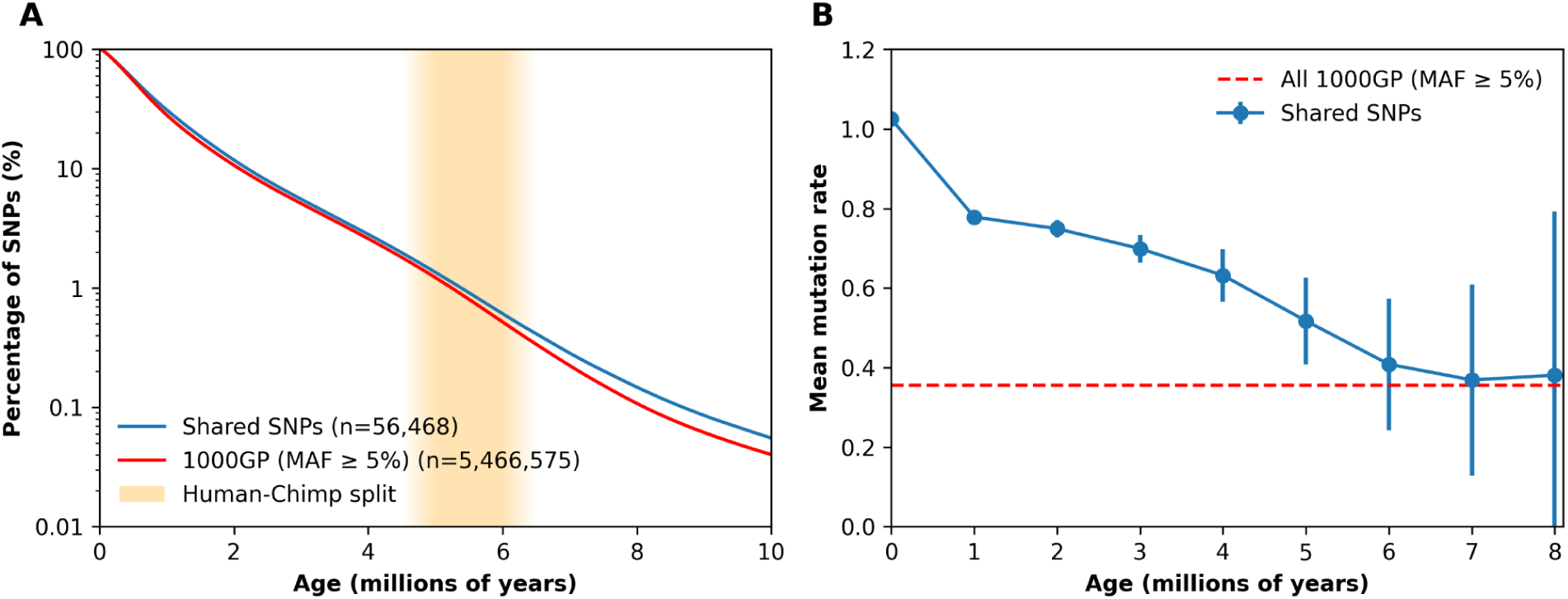
Common SNPs shared between humans and chimpanzees. (A) In blue are estimated ages of SNPs shared between humans and chimpanzees (n = 56,468) and and in red that of genome-wide common variants (1000 Genomes Project, MAF ≥ 5%; n = 5,466,575), plotted as the percentage of SNPs estimated to be older than a given age (on a log scale). The shaded band marks the human and chimpanzee species are thought to have split off from one another (∼5–6 Mya). (B) In blue is the mean mutation rate to the alternative allele across shared SNPs, for SNPs estimated to be older than the value on the x-axis; points show the mean and error bars the 95% confidence intervals. The dashed red line indicates the mean mutation rate to all 1000GP alternative alleles at ≥5% MAF. The mutation rates used are for the specific mutation type and context [56]. We note that the mean mutation rate shown for the 1000GP is higher than the genome-wide average mutation rate because the sites are ascertained to be common polymorphisms.

By this approach, it is clear that, as expected, the vast majority of shared SNPs are far too young to have been inherited from the human–chimpanzee common ancestor (Figure 1A and Supplementary Figure 2) and instead arose from recurrent mutation. Accordingly, shared SNPs are strongly enriched for mutation types that are highly mutable: as an illustration, the mean mutation rate across all shared SNPs is ∼2.9-fold that of common human variants (using per bp mutation rate estimates for humans [56]). This enrichment is driven in large part by CpG transitions, which make up more than half (55.6%) of shared SNPs but only 14.1% of common human variants. The mean mutation rate falls with increasing allele age (Fig 1B), as expected if older shared variants are more likely to reflect ancestral IBD sharing than recurrent mutations. Notably, by 6 Mya, which likely predates the human-chimpanzee split [45], the mean mutation rate of shared SNPs is indistinguishable from what is expected of common variants in humans, i.e., shows no evidence of enrichment of recurrent mutations.

### Genealogical reconstruction at putatively old shared SNPs

To identify ancient balanced polymorphisms, we focus on the subset old enough to plausibly represent variation shared IBD since the human–chimpanzee common ancestor. To that end, we select candidates by their estimated allele ages. Unlike for the age distributions in Figure 1A, here, we use a point estimate for each SNP. Specifically, we base ourselves on the midpoint age estimated by Relate and require it to exceed 4 Mya.

We deliberately set this threshold below the estimated human-chimpanzee split (5–6 Mya; [45,57]), because age estimates for old variants are subject to two downward biases. First, *Relate* underestimates the TMRCA in genomic regions with particularly ancient genealogies [54]. Second, summarising an allele by its branch midpoint introduces an additional bias that stems from the branching topology of coalescent trees. While the expected age of a neutral mutation given a branch assignment is the midpoint of the branch, once we condition on mutations that are truly old, that is no longer true. Mutations will fall in the older half of the deepest branches and using the midpoint will be a systematic underestimate; this effect may be big, since the deepest branches of a genealogy tend to be the longest [55] (Supplementary Figure 3). For the ancient variants of interest to us, the two biases compound, toward younger apparent ages. In this regard, we note that even our choice of 4 Mya may exclude some shared SNPs old enough to be IBD.

After excluding the MHC region, for which long-term balancing selection is already well established, we retain 860 shared SNPs (see Methods). For this set, we run *SINGER* on a combined human–chimpanzee sample. Tree reconstruction previously required an arbitrary choice of window size over which to estimate the local tree; if the window size chosen was too large, the signal could be missed [36]. ARG reconstruction methods overcome this limitation by inferring the location of changes in local tree structure due to ancestral recombination along the genome. To implement *SINGER*, we run three independent MCMC chains for each genomic region, and draw 100 samples from the posterior distribution of genealogies from each. Pooling the 300 sampled ARGs allows us to quantify the statistical support for a given topology and to test whether the trees around the focal SNP support it being IBD across species, i.e., lineages clustering by allele rather than by species (Supplementary Figure 4).

Of the 860 candidate SNPs, we are able to reconstruct local genealogies at 802 (93.3%, see Methods), of which 117 meet our TSP topology criteria in >80% of sampled ARGs and 70 in >90% (Supplementary Table 1). These 117 SNPs fall into 59 distinct genomic regions (see Methods; Supplementary Table 2), with 15 regions containing more than one TSP, spanning 173 bp–6.6 kb (Table 1).

**Table 1:**
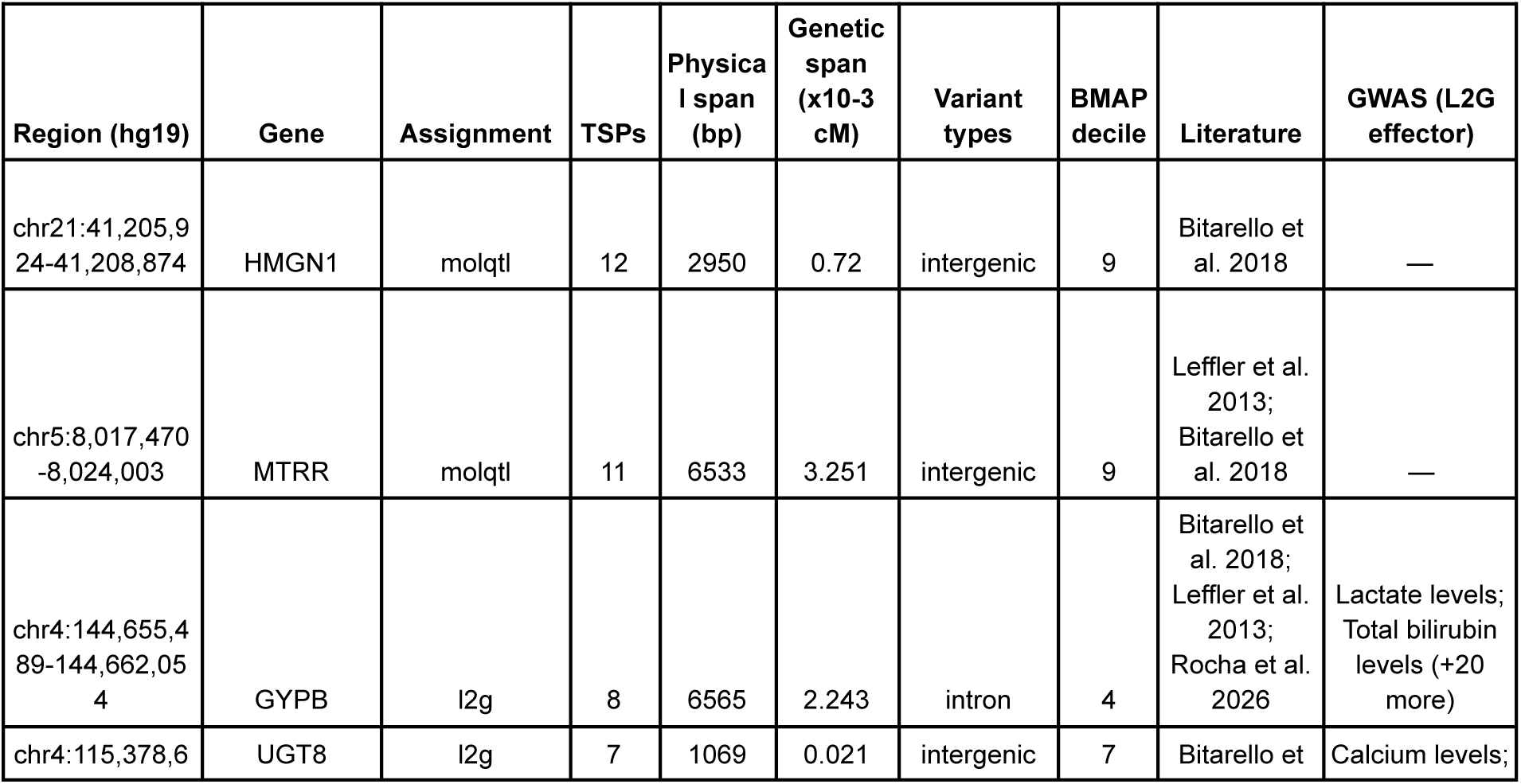
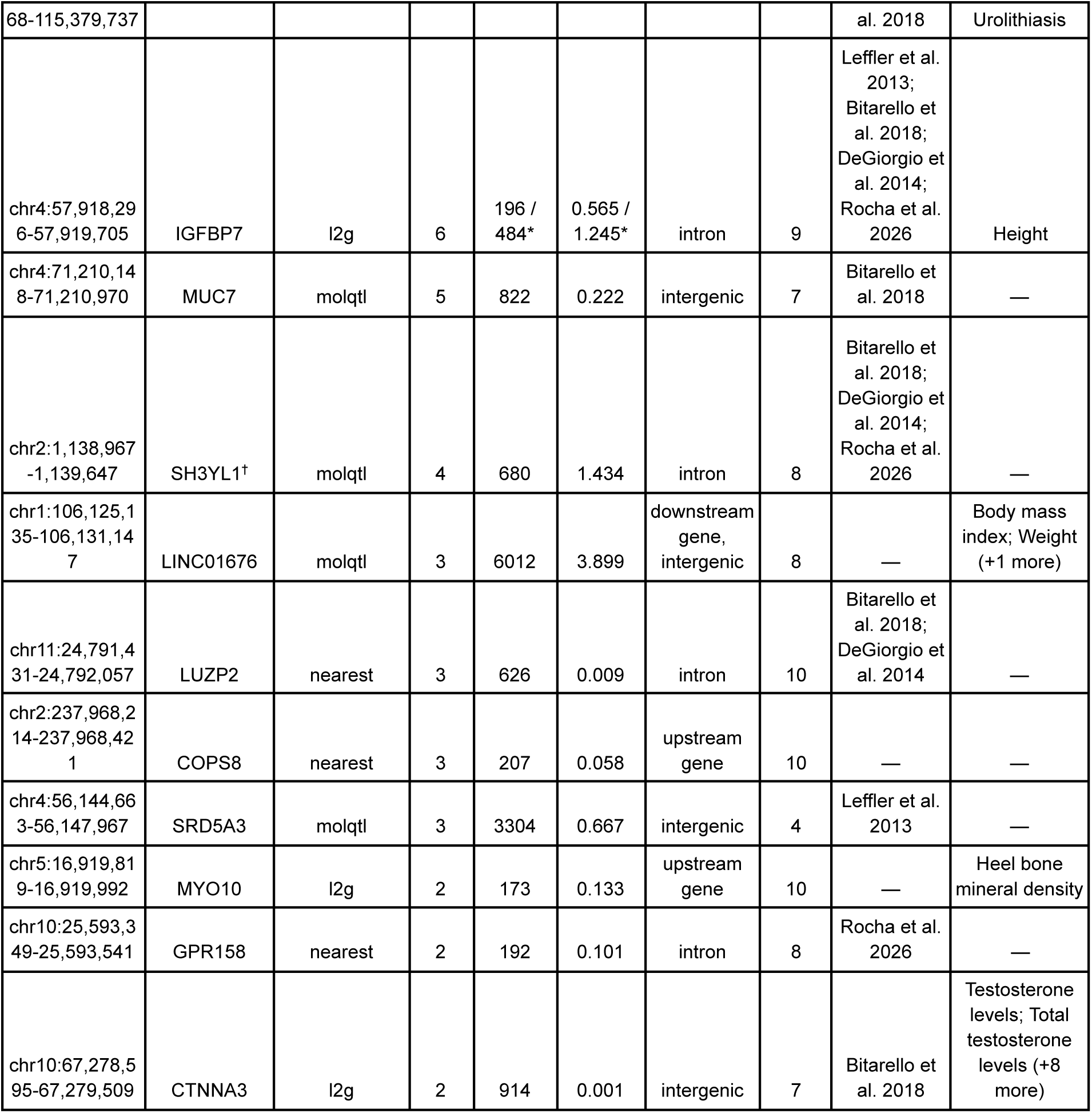
Regions with multiple trans-species polymorphic SNPs. The 15 regions in which more than one TSP was identified, ordered by the number of TSPs in the region. The column labeled “region” gives the genomic coordinates (hg19) of each cluster of TSPs, with SNPs within 10 kb assigned to the same region. “Gene” refers to the gene that the region is assigned to based on supporting evidence tiers (see Methods); “Assignment” details which of these tiers we use for the assignment. “TSP count” gives the number of shared SNPs in this region that are estimated to be ≥4Mya and which meet our TSP topology criterion in >80% of ARG samples. “Physical span” gives the maximum distance between genomic coordinates of TSPs in the region, and “Genetic span” provides the genetic distance, based on HapMap II [95]. “Annotation” refers to the SnpEff consequence class of the TSPs within the region. “BMAP decile” refers to the decile of the background selection map in which the region lies (1 = strongest, 10 = weakest background selection). “Literature” indicates prior studies that report the region as a long-term balancing-selection or TSP candidate. “GWAS (L2G effector)” lists genome-wide association study (GWAS) traits for which a TSP in the region is the index variant of a GWAS credible set and the assigned gene is the Open Targets locus-to-gene effector. **\***The IGFBP7 region is composed of two linkage groups (with 2 and 4 SNPs respectively); we report the spans of each of these linkage groups separately. **†** We assign the TSPs in this region to the gene *SH3YL1* based on molecular QTL evidence but note that the TSPs lie within the gene *SNTG2* which is reported by DeGiorgio et al. (2014) and Rocha et al. (2026) [28,58].

### Comparison to previously published results

Among the trans-species polymorphic regions that we identify, several have been reported elsewhere, either as TSPs or as candidates in scans for balancing selection in humans (Table 1 and Supplementary Table 2). The most comparable study to date is the scan for trans-species polymorphism conducted by Leffler et al. (2013) [38], who analyzed 59 humans and 10 chimpanzees sequenced to medium coverage, reporting 125 regions with multiple shared SNPs, of which six were highlighted as strong candidates. Three of those six also meet all of our criteria: those assigned by Leffler et al. to genes *FREM3* (assigned to *GYPB* in the present study; see below)*, IGFBP7* and *MTRR*. The other three are filtered out by our approach and may be artifacts: all variants assigned to *HUS1* and two of the three at *PROKR2* violate rules of Mendelian segregation in the chimpanzee pedigrees and the remaining *PROKR2* variant is estimated to be only ∼0.6 My old. In turn, the *ST3GAL1* variants fall within a segmentally duplicated region of the latest chimpanzee reference. Several of the TSPs we identify are also discussed in a preprint by Rocha et al. [58]. The authors reconstructed ancestral recombination graphs from physically-phased haplotype assemblies, including 58 chimpanzee and bonobo haplotypes together with 218 human haplotypes, and identified genomic windows in which coalescence times exceed 6 Mya in each species separately. Among the genes that they reported as having particularly old coalescence times in both humans and chimpanzee populations, three contain our TSPs: *IGFBP7, GPR158*, and *SNTG2* (as described below, we assign the TSPs within *SNTG2* to a different gene, *SH3YL1*, based on functional evidence). Additionally, the *FREM3/GYPB* locus is among their candidates and is the subject of a dedicated analysis in their study; we discuss this region further below.

We also compare our findings to scans for balancing selection carried out using human variation data alone. These studies relied on summaries of the polymorphism data; since such statistics reflect properties of the underlying ARG, findings based on them do not provide independent corroboration of ours. Nonetheless, overlap among studies indicates that multiple aspects of the underlying tree fit the expectations of a balanced polymorphism, rather than only the features relied on here to identify TSPs. Notably, several of the TSP regions overlap with the top-ranked genes reported by DeGiorgio et al. [28]. To identify the genomic signature of long-term balancing selection, the authors used two composite likelihood ratio statistics that test for an excess of polymorphism relative to divergence and a shift toward intermediate allele frequencies expected at neutral sites linked to a balanced polymorphism. The statistical evidence was evaluated for each site, and genes were ranked by the maximum score across sites between transcription start and stop. We consider a region as identified in both our study and theirs when a TSP falls within the 141 genes that they highlight as of interest. The overlap again includes *IGFBP7* and *SNTG2,* as well as *LUZP2*, *RNF150* and *RBFOX1,* which have not previously been reported as shared with chimpanzees (though Rocha et al. also report deep coalescence within chimpanzees at *LUZP2* and *RBFOX1* [58]). One of these loci, *RNF150*, was also identified by Rasmussen et al. [34], who inferred ancestral recombination graphs genome-wide in humans and ranked 10 kb windows by their posterior mean TMRCA. The TSP in this gene falls within one of the twenty deepest windows in the genome, which they estimated to be ∼325K generations old (approximately 9.4 Mya for a generation time of 29 years). Finally, a larger set of our regions overlaps some of the candidate windows identified by Bitarello et al. [32]. Their approach was to look for shifts in minor allele frequencies consistent with long-lived overdominance and to pinpoint candidate windows whose values are furthest from expectation in simulations of neutrality. We count a region as overlapping where it intersects one of the windows that they defined as outliers (top 0.05% of the genome-wide distribution), in any of their 12 scans. Sixteen of our 59 regions do so, including nine of the 15 regions harboring more than one TSP. Thus, in total, approximately a third of the 59 regions that we identify as harboring a TSP have been previously reported as under long-lived balancing selection, either based on patterns in humans alone or by an analysis of humans and chimpanzees.

### Uncoalesced regions in humans and chimpanzees

True trans-species polymorphisms must fall in regions of the genome where at least two lineages remain uncoalesced in humans and in chimpanzees at the time of their split. We therefore take a second approach to identify such regions by inferring local coalescence times under the standard PSMC model [59] and extracting, via the posterior-decoding step implemented in *cobraa* [60], the posterior probability that the two haplotypes of each diploid individual remain uncoalesced >5.5 Mya (in each 1 kb segment of the genome). We focus on this timepoint because it roughly aligns with the peak of the coincident increases in the effective population sizes of humans and chimpanzees inferred by Cousins et al. [45], which the authors interpret as evidence of some ongoing gene flow between incipient species. Applying this method to our chimpanzee samples and combining the results with the published human estimates of Cousins et al., we sum these per-individual probabilities within each segment, separately per species. We retain segments whose summed probability exceeds 1, such that in expectation at least one sampled individual carries an uncoalesced haplotype pair.

Intersecting the segments passing this criterion in humans and in chimpanzees identifies a set of regions inferred to have remained uncoalesced beyond 5.5 Mya in both species. In total, these regions cover 0.23% of the human autosomal genome that we analyze. As expected, our putative TSPs are highly enriched within these “reciprocally uncoalesced regions”: whereas only 0.7% of 1000GP common SNPs and 1.5% of all shared SNPs lie in such regions, 16.6% of shared SNPs whose estimated age is ≥4 Mya do. For the set of SNPs that we infer to be TSPs, the fraction rises to 47.0% (Figure 2). In this regard, we note that the overlap is not expected to reach 100%, because the set of uncoalesced regions that we identify is not exhaustive. Trivially, because we look for uncoalesced segments in nine chimpanzee individuals and seven humans rather than the full sample for which we infer TSPs, so deep lineages absent from these individuals will go undetected. But also because the approach infers uncoalesced lineages only within a single diploid genome and therefore will miss cases in which there are two deeply divergent lineages in the sample, but no current day individual is carrying both.

**Figure 2:**
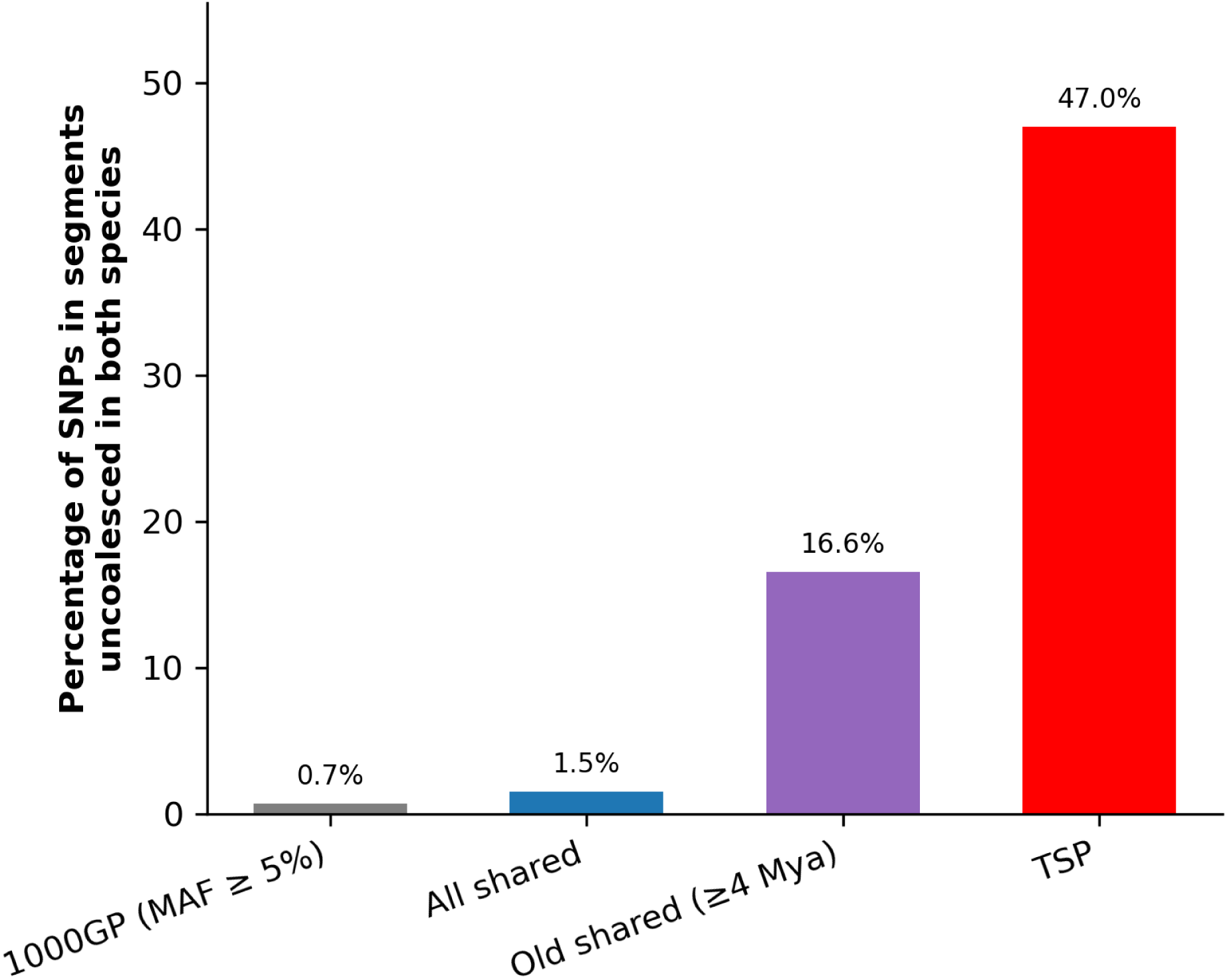
Enrichment of trans-species polymorphisms within segments uncoalesced in both species by 5.5 Mya. Shown is the percentage of SNPs in each set that fall within segments that are uncoalesced by 5.5 Mya in both humans and chimpanzees, for all common variants in humans (in grey; 1000 Genomes Project, MAF ≥ 5%); all human-chimpanzee shared SNPs (in blue), old shared SNPs (in purple; defined as having an allele age estimate ≥ 4 Mya) and trans-species polymorphisms (in red).

### Evidence that trans-species polymorphisms between humans and chimps are largely neutral

TSPs between humans and chimpanzees have long been treated as the gold standard for evidence of long-term balancing selection [4,36]. Under neutrality, a TSP requires at least two lineages from each species to remain uncoalesced until at least the time at which the species split. In any one species, the probability of two lineages not having coalesced decays exponentially with the split time (relative to effective population size, that is at rate e^-T/2Nₑ^ per lineage) [36]. Because the human-chimpanzee split is old relative to widely-used estimates of Nₑ in humans, this probability was considered to be negligible, and on that basis, a polymorphism shared IBD since the common ancestor was taken as compelling evidence for selection [36]. However, the findings by Cousins et al. [45], which indicate that a non-negligible fraction of the human genome remains uncoalesced beyond ∼5 Mya, raise the possibility that trans-species polymorphisms occur by chance (i.e., under neutrality) at an appreciable rate.

If the TSPs that we identify are under balancing selection, we expect them to have a functional effect, altering protein-coding or regulatory sequence. Therefore, we would predict that the set of TSPs will be enriched for functional consequences, and the genes with which they are associated might share a common biological role or pathway. If instead most TSPs are neutral, they will simply fall in the regions of the genome most likely to remain uncoalesced by chance, and are not expected to carry a functional signature beyond that of the shared variation from which they are drawn. We evaluate these expectations, then look more broadly at the regions of the genome that remain uncoalesced at 5.5 Mya and their properties.

As a starting point, we ask whether, as a class, the TSPs are enriched for variants under selection. We first do so by considering their annotations, loosely ordered by their typical fitness consequences (Figure 3A). We find that TSPs are overwhelmingly non-coding: 60.7% fall in intergenic sequence and 28.2% in introns (88.9% combined), with a further 11.1% in gene-flanking regions (within 5 kb of a gene); none of the 117 TSPs directly alter the protein-coding sequence (Figure 3A). Moreover, the TSP variant effects are shifted toward less consequential fitness effects relative both to genome-wide common variation (1000GP, MAF ≥ 5%) and to all shared SNPs: the fraction of TSPs that is intergenic is greater than in either of these sets, whereas the fraction in sequences flanking genes is lower (Figure 3A). Similarly, considering their overlap with candidate cis-regulatory elements (cCREs), we find that 15.4% (18/117) of TSPs overlap a SCREEN cCRE, which is lower than the 22.3% of common 1000GP variants and 20.7% of all shared SNPs. There is therefore no evidence for an enrichment of functionally-important annotations among TSPs.

**Figure 3:**
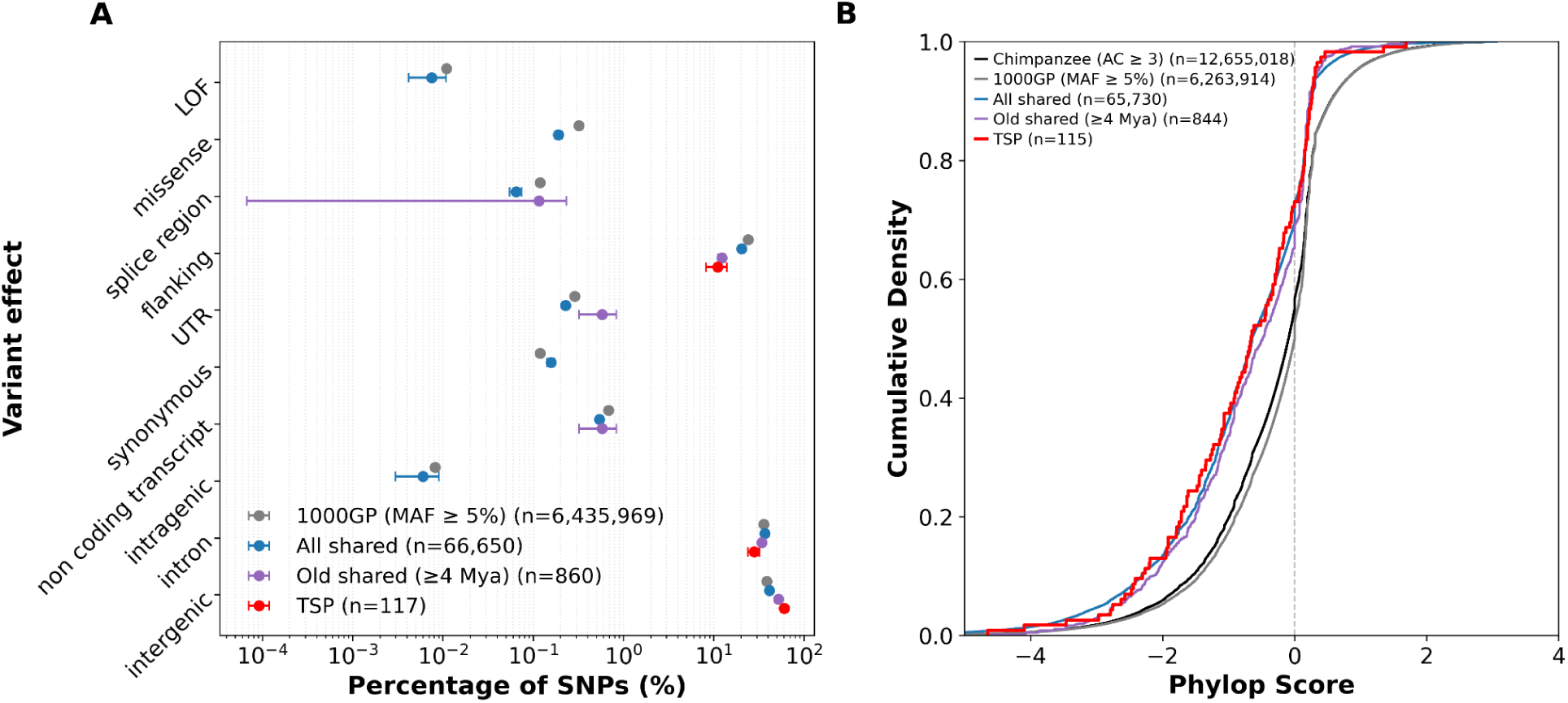
Functional characterization of trans-species polymorphisms. (A) Distribution of predicted variant effects across four SNP sets: all common variants in humans (in grey; 1000 Genomes Project, MAF ≥ 5%), all human-chimpanzee shared SNPs (in blue), old shared SNPs (in purple, defined as having an estimated allele age ≥ 4 Mya) and trans-species polymorphisms (in red). Each SNP was assigned its single most severe consequence across overlapping transcripts annotated by SnpEff (see Methods). The UTR category includes 5’ and 3’ UTR variants are merged and “flanking” variants within 5Kb of the transcription start site or transcription end site. Categories are loosely ordered by their typical fitness consequences (highest at top), based on [82]. Points denote the percentage of SNPs per category, shown on a log scale; error bars show the binomial standard error. The total number of SNPs in each set, n, are given in the legend. (B) Cumulative density function of phyloP conservation scores for the four SNP sets shown in panel A (same colors as in A), as well as for common chimpanzee variants (in black; allele count of at least three in 43 unrelated chimpanzees).The total number of SNPs in each set with an available phyloP score, n, are given in the legend. phyloP scores are taken from Fu et al. (2014), and were computed on the Ensemble Compara 36-eutherian-mammal EPO alignment after masking out the human sequence.

Next, we consider levels of constraint at TSP sites relative to other sites in the genome by comparing phyloP conservation scores across SNP sets (Figure 3B; see Methods). PhyloP scores are based on multiple-species alignments, which include a single reference sequence for each species. To avoid the TSP itself having an effect on the phyloP score, we rely on phyloP scores computed from a 36 eutherian mammal alignment in which the human sequence is masked [61]. The chimpanzee genome, however, remains in the alignment, potentially leading to a decreased PhyloP score whenever the chimpanzee reference happens to carry the derived allele, as is more likely to happen for TSPs. To assess how large this bias may be, we also examine the distribution of scores for sites that are variable within the chimpanzee sample (with an allele count ≥ 3, i.e., the same threshold applied to the chimpanzee side of the shared set), as this set should suffer from the same problem. Overall, TSPs show no evidence of elevated constraint: their scores lie below those of common chimpanzee variants (median −0.66 versus -0.09; Kolmogorov–Smirnov D = 0.22, p = 3.5×10⁻^5^), and the distribution of TSP scores is indistinguishable both from shared SNPs as a whole (p = 0.96) and from the ≥4 Mya shared subset (p = 0.55). A caveat to these results is that if our TSPs are subject to balancing selection in other species across the mammalian tree, it may shift phyloP scores towards unconstrained values.

If TSPs are neutral, we might expect them to be preferentially found in regions with less effect of linked selection. Notably, background selection, which is strongest in gene-dense, low recombination regions, increases coalescence rates (i.e., decreases the effective population size) and thus decreases the probability that lineages remain uncoalesced until the split time [62,63]. To evaluate the genomic distribution of TSPs in this regard, we use previously generated estimates of the strength of background selection (BMAP) for human autosomes [64]. The BMAP score quantifies the extent of the reduction in the local effective population size due to background selection [64]. As expected, when considering genomes from different species and subspecies, the proportion that remains uncoalesced > 5.5 Mya reflects the diversity levels within a genome (Supplementary Figure 5). Also as expected, within each individual genome, this proportion increases monotonically as background selection weakens (Supplementary Figure 5). This pattern becomes more pronounced when looking at the proportion of the genome that remains uncoalesced in both species: it rises ∼27-fold across the deciles, from 0.024% in the first to 0.626% in the tenth (Figure 4A). Turning to the TSPs, their number mirrors this gradient, consistent with what would be expected under neutrality (Figure 4B).

**Figure 4:**
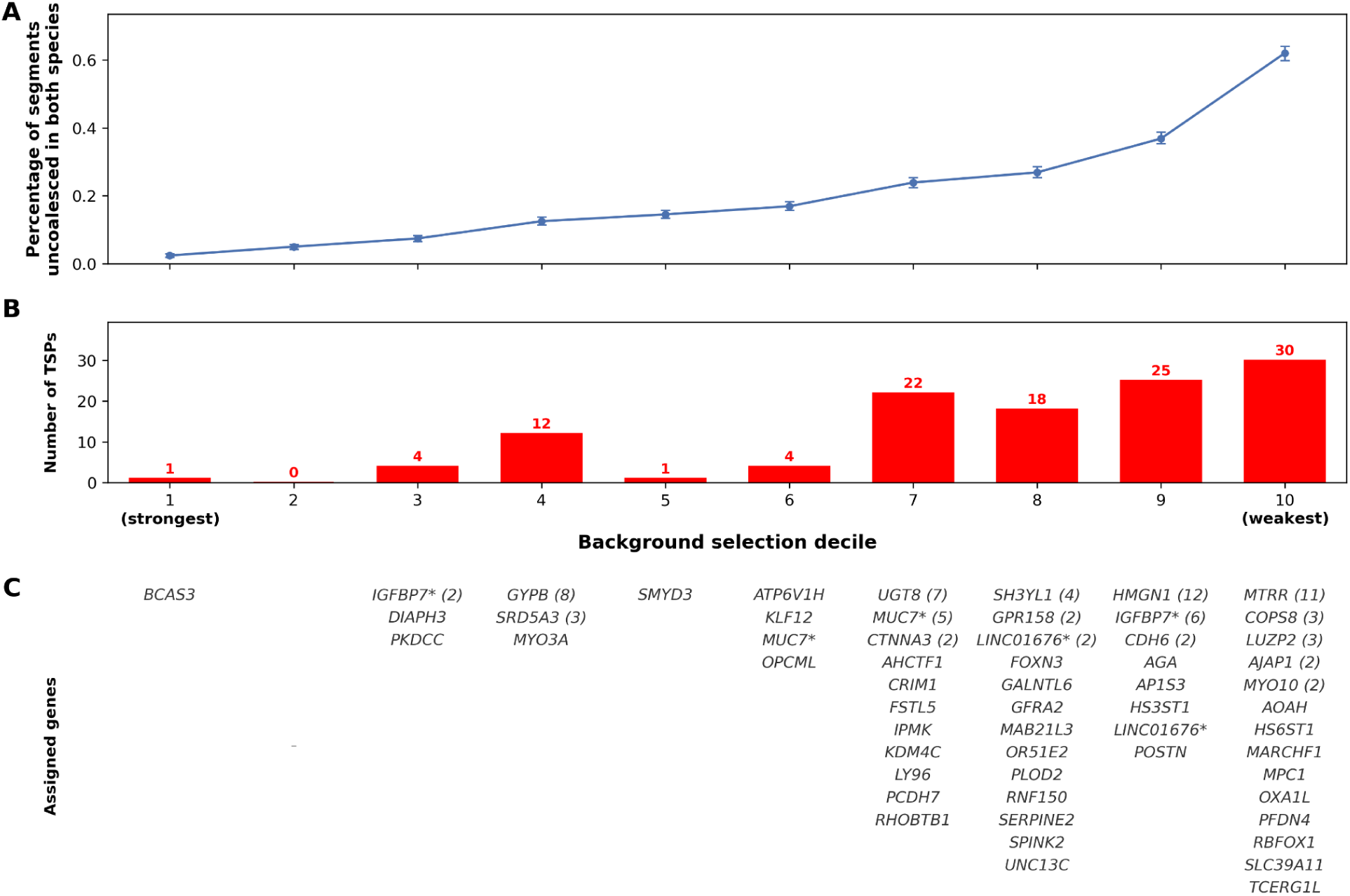
Trans-species polymorphisms and segments uncoalesced in both species by 5.5 Mya, stratified by strength of background selection. The autosomal genome (MHC excluded) was partitioned into deciles of background selection scores (BMAP). (**A**) Shown is the percentage of sequence that is uncoalesced in both humans and chimpanzees by ∼5.5 Mya for each decile, i.e., the fraction of sequence callable in both species that is uncoalesced in both species (see Methods). Points show the percentage per decile and error bars 95% bootstrap confidence intervals. (**B**) Reported is the number of TSPs (from a total of 117) in each decile, both with the bar height and number. (**C**) Listed are the genes to which the TSPs are assigned in each decile. An asterisk marks the genes that are assigned TSPs in more than one decile.

As a complementary approach, we assess whether the genes linked to the TSPs are enriched for particular biological functions, as might be expected if the polymorphisms were maintained by selection related to a common functional role. Previous studies reported cell-surface glycoproteins to be enriched for variants shared between humans and chimpanzees [38]. Furthermore, the best-established trans-species polymorphisms fall in similar categories: MHC products are cell-surface glycoproteins and ABO encodes a glycotransferase that determines cell-surface glycan antigens. To test whether a related signal is seen among our TSPs, we assign each of the 59 TSP regions to a single gene using a tiered rule, based on the quality of functional evidence linking variants to genes, yielding 55 distinct genes (see Methods). A naive comparison against all autosomal protein-coding genes suggests an enrichment of glycoproteins: 44% (24/55) of the TSP genes are in this category, versus 22% genome-wide (1.9-fold; hypergeometric p = 0.0004); membrane glycoproteins specifically show a weaker, non-significant enrichment: 22% (13/55) versus 17% genome-wide (1.4-fold; hypergeometric p = 0.11).

As shown by our previous analyses, however, TSP loci are not a random sample of the genome: they tend to be found in gene-sparse, highly recombining regions, which are themselves atypical in gene content. To control for these features, we compare the set of genes with associated TSPs against 2,000 sets of randomly-chosen genes in regions matched for BMAP scores and SNP numbers (see Methods). Against this matched background set, the glycoprotein enrichment is no longer significant (1.3-fold; empirical p = 0.06) and membrane glycoproteins are no longer enriched at all (0.9-fold; empirical p = 0.68). These findings suggest that the apparent signal is an artefact of where TSPs are found in the genome.

Extending this approach to consider Gene Ontology, KEGG, Reactome and MSigDB Hallmark categories, we test the 44 categories containing at least three TSP genes against the matched null. Ten are significant at a Benjamini-Hochberg FDR of 0.05, collapsing to six groups once categories with substantial overlaps in genes are combined. The largest group relates to the endomembrane system (lysosome, lysosomal and lytic vacuole membrane, trans-golgi network and endosome membrane), and is driven by genes *AGA*, *AP1S3*, *ATP6V1H*, *MARCHF1*, *POSTN* and *RBFOX1.* The other significant categories are endosome membrane (*LY96*, *MARCHF1*, *OR51E2*, *RHOBTB1*), oxidative phosphorylation (*ATP6V1H*, *MPC1*, *MTRR*, *OXA1L*; 9.9-fold, FDR-adjusted p = 0.015), regulation of filopodium assembly (*BCAS3*, *MYO10*, *MYO3A*; 12-fold), RHO GTPase effectors (*AHCTF1*, *DIAPH3*, *KDM4C*) and epithelial-mesenchymal transition (*CDH6*, *PLOD2*, *POSTN*, *SERPINE2*). Each of these individual categories rests on three or four genes, so individually they are tentative and together they account for a minority of the TSP genes (21 of 55). Indeed, for the 15 regions with multiple TSPs, about which we are most confident, only three belong to these categories. Thus, once genomic features of TSP regions are taken into account, there does not appear to be a clear common function shared among a large proportion of TSPs.

### Evidence that not all TSPs are neutral

These analyses reveal no evidence that as a set, human-chimpanzee TSPs are enriched for specific functions or unusually strong targets of selection, but they do not establish that every TSP is neutral. Indeed, for a subset of loci, there is independent evidence of function that suggests the locus may be the target of ancient balancing selection.

The clearest is within the glycophorin (GYP) locus on chromosome 4, in which we find eight tightly linked TSPs spanning 6,565 bp. Trans-species polymorphisms at this locus were first reported by Leffler et al. (2013) and have subsequently been the subject of further investigation, owing to the discovery of a nearby, linked locus implicated in malaria resistance [65]. The genes within this locus, GYPA and its great-ape specific duplicates GYPB and GYPE, are erythrocyte receptors for *Plasmodium falciparum* invasion; the GYPA–GYPB (Dantu) fusion reduces the risk of severe malaria by up to ∼74% [66]. In a recent pangenome analysis of the *Pan* genus, Rocha et al. 2026 [58] reported elevated TMRCAs at this locus in both chimpanzees and humans and extensive recurrent structural evolution, including multiple independent Dantu-type GYPA–GYPB fusions in humans and six novel chimpanzee glycophorin genes formed by ancient fusion and interlocus gene conversion. Our gene-assignment procedure assigns the TSPs that we identify to GYPB on the basis of the Open Targets GWAS locus-to-gene score (0.83, narrowly above FREM3 at 0.79), but the same variants also fall within eQTLs for the neighbouring glycophorin gene GYPE.

Beyond this locus, we identify a number of new candidates for balancing selection. Notably, in the secretory calcium-binding phosphoprotein (SCPP) gene cluster, there are five linked TSPs spanning 822 bps in the intergenic region between *CABS1* and *SMR3A* (Figure 5). This cluster of genes, which includes salivary proteins as well as genes with roles in tooth mineralization and calcium homeostasis in milk [67], is a known hotspot of evolution in mammals, with extensive gene diversification, including great-ape specific genes [68]. The TSPs in this region are tightly associated with one another and are found at frequencies of around 40-50% in all human continental groups; in chimpanzees, the average frequencies across chimpanzee subspecies are similar but the derived allele is fixed in the Western sample (Figure 5C). While these SNPs are not within a known regulatory element, they are among the credible set of variants for an expression quantitative trait locus (eQTL) that affects expression of *Mucin 7* (*MUC7*) in skin [69].

**Figure 5:**
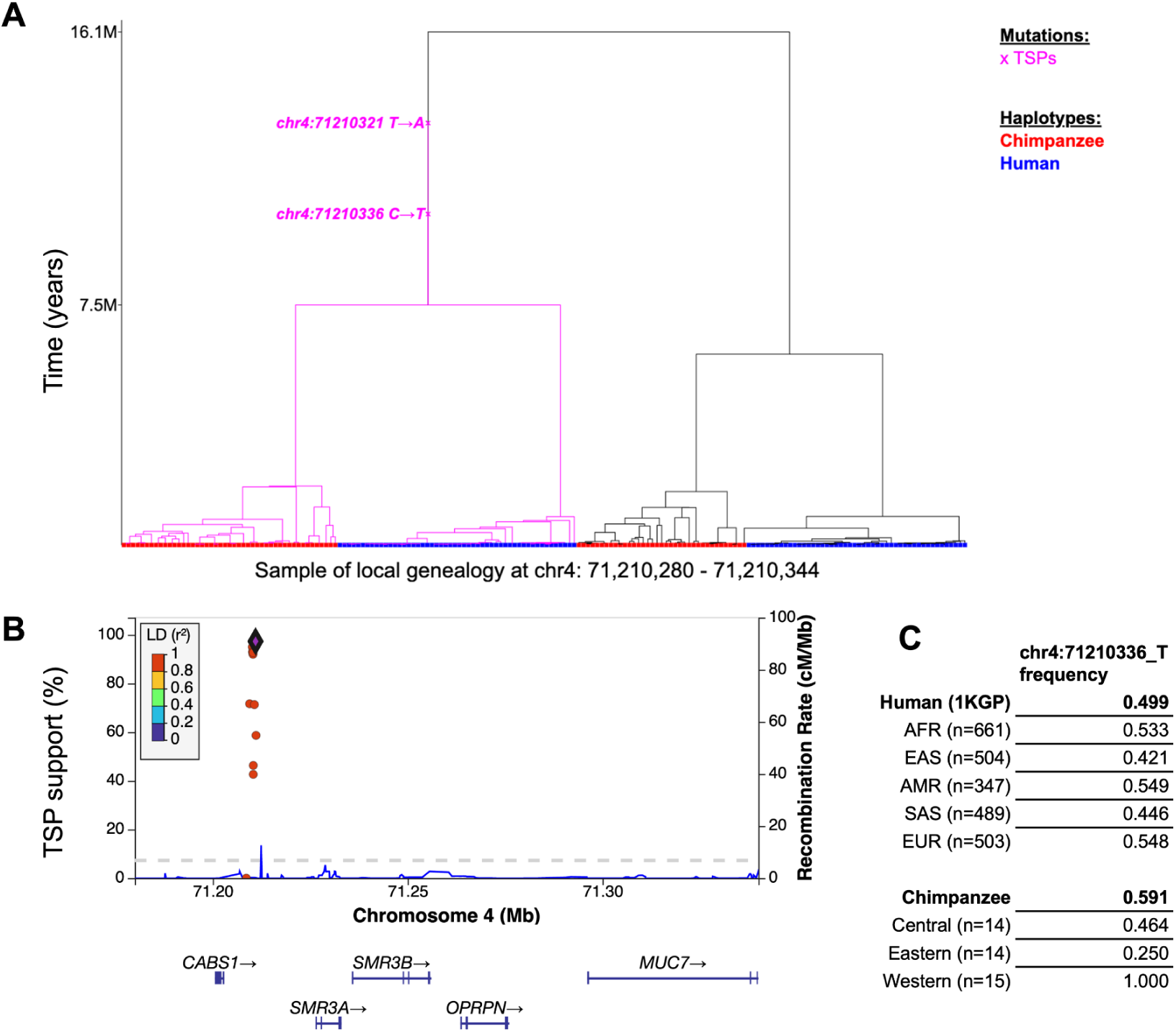
Example of a novel trans-species polymorphism associated with *MUC7*. (**A)** Local genealogical reconstruction using *SINGER* on the combined sample of 50 human and 43 chimpanzee samples for the region chr4:71,210,280 - 71,210,344 (GRCh37/hg19 coordinates), which lies in BMAP decile 7 (see Figure 4). Checkmarks denote mutations, branches below the TSP mutations are colored pink, and the coloring at the tips of branches indicates whether the haplotype is from a human (in blue) or a chimpanzee (in red). See Methods for details. (**B**) LocusZoom plot of old shared SNPs in this region. On the x-axis is the genomic position; on the y-axis to the left, the proportion of samples from the posterior distribution of *SINGER* tree topologies that support the old shared SNP (defined as having a *Relate* YRI estimated allele age ≥ 4 Mya [47]) being shared identical by descent (“TSP support”); and on the y-axis to the right is the recombination rate along the chromosome based on Hapmap II [95]. Points representing each shared SNP are coloured by r2 value with chr4:71210970_A, which has the highest TSP support (meeting our criteria in 97.3% of trees; see Methods). Nearby protein-coding gene locations are shown below; the TSPs lie between genes *CABS1* and *SMR3A* but are fine-mapped to an eQTL affecting expression of *MUC7* in the skin (see main text). (**C**) Allele frequencies for chr4:71210336_T, one of the five TSPs in this region, in different population and subspecies samples.

*MUC7* is a secreted, antimicrobial protein thought to play an important immune role in clearing bacteria and other microbes from the oral cavity, a function mediated by densely O-glycosylated repeat domains that bind microbial surfaces [70]. In the minor salivary gland, the only salivary tissue for which eQTL data exist, the TSPs show no association with MUC7 expression (p = 0.20, n = 180; GTEx v10), despite MUC7 being highly expressed; whether there is an effect in the major salivary glands, which secrete most salivary *MUC7*, is to our knowledge unknown. *MUC7* has an unusually dynamic evolutionary history in primates: its antifungal histatin-like domain shows evidence of positive selection on the primate lineage and its copy-number variable subexonic repeats have evolved recurrently across primates [71] and within the human lineage, where a deeply divergent haplotype in African populations has been attributed to introgression from an unidentified archaic hominin [72]. The TSPs that we identify lie close to, but are distinct from and not in LD with, that introgressed haplotype. Given that the high diversity at *MUC7* has been interpreted as a response to diverse pathogen pressures [71], and balancing selection is a plausible outcome of such pressure where no single variant is universally advantageous, this locus is a promising candidate for further investigation.

A second intriguing example is *SRD5A3*, near which we find three linked TSPs within 3,304 bps. This locus was reported among the shared-polymorphism regions of Leffler et al. (2013) but was not one of the six loci that were resequenced in order to reconstruct a local genealogy; our reconstruction provides that confirmation (Table 1). Two features make it a strong candidate for ancient balancing selection. First, unlike most TSPs, which are in regions of low background selection and thus are more readily explicable by chance, the *SRD5A3* TSPs lie in BMAP decile 4. Second, the three TSPs form a single tightly-linked haplotype (r² = 0.94–1.0) that tags a *SRD5A3* transcript-usage QTL in thyroid (GTEx; β = +0.41, P = 3×10⁻⁶), with near perfect LD with the credible-set lead variant in Europeans (r² = 0.97–1.0). *SRD5A3* (steroid 5α-reductase 3 / polyprenol reductase) catalyses an essential step in dolichol synthesis and protein N-glycosylation, and its loss causes a congenital disorder of glycosylation, so a common variant shifting *SRD5A3* isoform usage is a plausible functional target. This locus too therefore seems like a promising candidate.

#### Selection pressures on regions with deep coalescences

Finally, we ask whether regions that are uncoalesced in the two species, whether or not they carry a detectable TSP by our approach, show evidence of being under balancing selection. In the absence of selection, the probability that a site would be uncoalesced in both species by the split time is the product of the probability that it did not coalesce in humans and did not coalesce in chimpanzees. If instead there is selection acting on these genomic regions, and these selection pressures are shared, then regions uncoalesced in one species will be more likely to be uncoalesced in another. We expect a correlation in the regions uncoalesced in both because of direct purifying selection on sites themselves and background selection, as well as potentially because similar balancing selection pressures maintain variation in the two species.

To account for direct and indirect purifying selection, we stratify the genome by the strength of background selection (considering quartiles of BMAP) and restrict the analysis to unconstrained sites (using phyloP; see Methods). We then consider whether the fraction of regions uncoalesced in both species exceeds what would be expected if it occurred independently in the two species. We find such an effect in every quartile and intriguingly, it is most pronounced where background selection is strongest (Figure 6). Moreover, if we add back the MHC region, which we excluded from previous analyses,, this signal of enrichment in the quartile of strongest background selection is further increased (Supplementary Figure 6). Thus, the regions with deep coalescences tend to overlap more than expected in humans and chimpanzees, even after our attempt to account for shared effects of background selection and direct constraint. While this finding could reflect shared selection pressures not captured by our controls, it is also consistent with shared balancing selection pressures maintaining ancient lineages at a subset of loci in both species.

**Figure 6:**
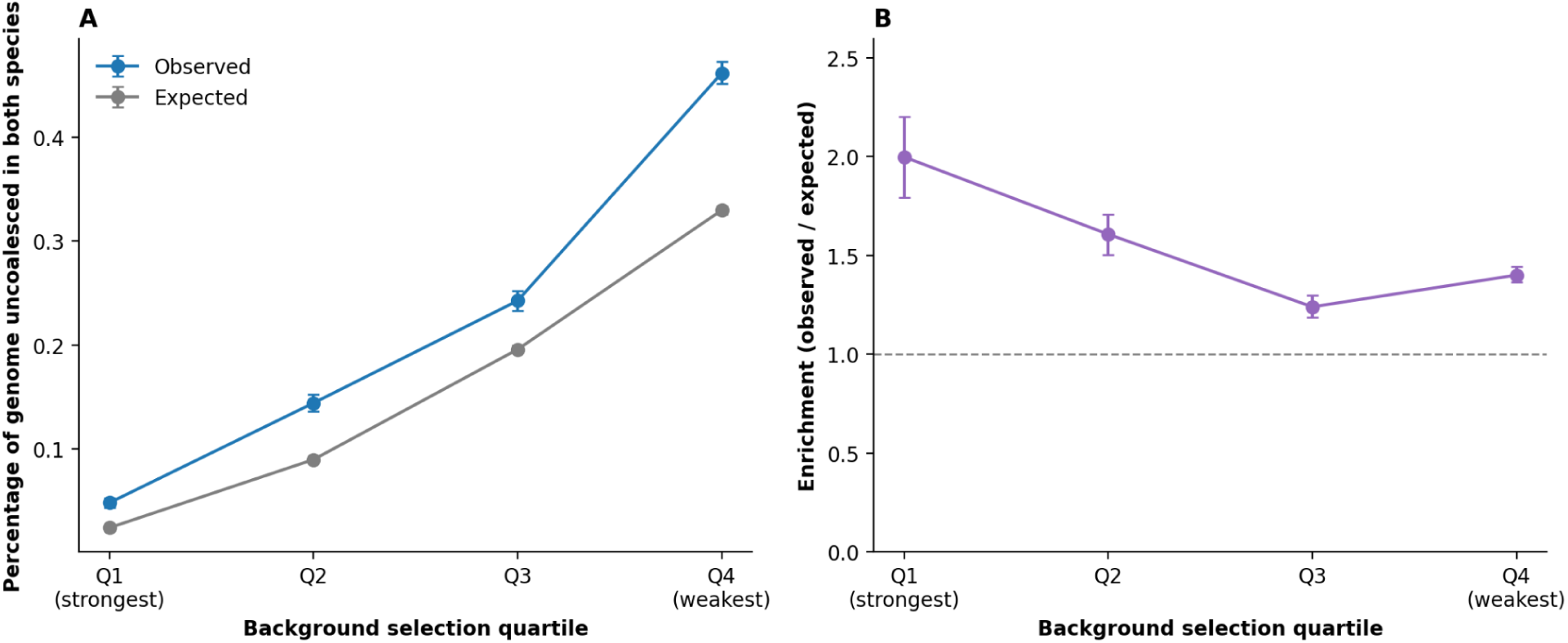
Regions uncoalesced in both species relative to what would be expected if independent, stratified by the strength of background selection. Autosomal regions were split into quartiles, based on the strength of background selection. Constrained segments were excluded, as was the MHC (see Methods). (**A**) For each quartile, we report the estimated proportion of the sequence that is uncoalesced in both humans and chimpanzees (in blue) by 5.5 Mya versus what we would expect assuming that coalescent events are independent in the two species (see Methods; in grey). (**B**) The ratio of observed to expected proportions, as a measure of enrichment. Points are estimates and error bars 95% bootstrap confidence intervals.

As is the case for the TSPs, a naive test suggests that the genes closest to regions uncoalesced in both species show an enrichment of membrane glycoproteins, in every background selection quartile (1.2 to 2.1 fold, hypergeometric p ≤ 1.7×10⁻³). Once we account for the tendency of long and more isolated genes to have larger genomic regions assigned to them, this enrichment disappears in all except the quartile with the strongest background selection. There, membrane glycoproteins remain two-fold enriched (p=0.005), whereas in the remaining quartiles, the fold-enrichment falls to 1 to 1.1-fold (p>0.14). This signal is independent from the TSPs identified, as none of the genes that drive the signal are among those assigned to our TSP regions.

This enrichment test also identifies olfactory receptors as enriched in the second and third quartiles (6.9-fold, p=0.024 and, 6.6-fold, p=0.014 respectively). The significance of this enrichment is unclear, however, as olfactory receptors are known to evolve under little selection in humans and to be clustered in subtelomeric regions of the genome [73,74]. One further case of interest is worth noting but tentative: in the quartile with the weakest background selection, genes involved in antifungal immunity are enriched 5.7-fold, driven by three C-type lectin receptors (*CLEC4C*, *CLEC6A*, *CLEC4D*) and *PLCG2*.

Taken together, there are therefore a handful of loci with independent functional information that are plausible candidates for ancient balancing selection, including the glycophorin locus, *MUC7*, *SRD5A3*, and *IGFBP7*, suggesting that a small number of trans-species polymorphisms were maintained by selection rather than having been maintained by chance. The finding that regions with deep coalescence times tend to be shared between the two species more than expected under neutrality or purifying selection also provides tentative evidence for the influence of balancing selection, as does the finding of an enrichment of membrane glycoproteins among regions that remain uncoalesced despite strong background selection.

## Discussion

By combining genome-wide sequencing of chimpanzees and humans with local ancestral recombination graph reconstruction, we systematically catalogue the single-nucleotide polymorphisms shared IBD between humans and chimpanzees, distinguishing genuine trans-species polymorphisms from the far larger set of recurrent mutations at mutable sites. This approach yields 117 TSPs in 59 regions, including both established balancing-selection loci and novel candidates, and shows that ARG-based methods can resolve the short (∼100 bp–1 kb) genealogical footprint expected around such ancient variants, a resolution unavailable to the window-based approaches used when these variants were last catalogued a decade ago [38].

Our central result is that, as a class, these trans-species polymorphisms show no clear signatures of selection. They are overwhelmingly non-coding and, if anything, depleted for functional and regulatory consequence relative to shared variation generally; they are no more constrained than a typical site; and once features of their genomic locations are taking into account, the set of genes to which they are associated shows no significant enrichment in gene function. As suggested by the recent finding that a non-negligible fraction of the genome remains uncoalesced by the human-chimpanzee split [45], our findings make clear that the existence of a shared, polymorphism that is IBD between humans and chimpanzees is not, in itself, sufficient evidence of long-term balancing selection. Instead, the great majority are most parsimoniously explained as neutral, incomplete lineage sorting. Indeed, we find that TSPs fall in the regions most prone to deep coalescence by chance, namely where background selection is weakest. Leffler et al. (2013) and Rasmussen et al. (2014) had previously estimated high false discovery rates for targets of long-lived balancing selection [34,38]. Our findings indicate that these rates may be yet higher.

One implication is that there may be little ability to distinguish real functional enrichment signals from spurious ones. In great apes, based on our most well-characterised examples (MHC and the glycophorin locus), as well as an enrichment observed among broader sets of trans-species polymorphisms [38,44], it has been argued that long-term balancing selection is driven by host–pathogen interactions, with cell-surface and membrane glycoproteins as its archetypal targets. While we also recover an apparent glycoprotein enrichment among genes to which TSPs are assigned, the signal dissipates when the background set is matched for salient genomic features. This result highlights the more general point that genes of a given function are not randomly distributed across the genome. There may well exist functions that are enriched among true targets of balancing selection. Our results do not argue against this possibility and indeed we see a hint of a signal for membrane glycoproteins in regions of the genome subject to strong background selection but that nonetheless have deep coalescent times (Figure 6). Rather, our results suggest that the false-positives contained within a set of candidate balanced polymorphisms may themselves cause artefactual enrichment signals. In this regard, we note that candidate loci ascertained through other methods, such as scans for genome-wide tMRCA outliers, should be subject to the same pitfalls.

These considerations do not mean that every TSP is neutral. For a minority of loci shared between humans and chimpanzees, there is independent evidence that suggests they are subject to selection, notably at the glycophorin locus, where the malaria-protective Dantu variant provides direct functional support [65]. For a few other loci, including the variants in an eQTL for the salivary mucin *MUC7* and the transcript-usage variants at *SRD5A3*, their genomic and functional features make them plausible candidates.

Furthermore, while our findings have implications for our understanding of many TSPs between humans and close evolutionary relatives such as chimpanzees, they do not call into question the interpretation of TSPs that have been established to be extremely old. Specific examples in primates, such as those at the ABO locus and the MHC, have been maintained over timescales that require balancing selection [35,37,73]. There may be additional balanced polymorphisms that are similarly old but essentially invisible to existing methods. Notably, if the target of selection is a single variant, then over a sufficiently long timescale, recombination will cause the length of the associated haplotype to decay to the point where accurate reconstruction of the local topology is impossible [36]. Thus it is possible that other extremely old polymorphisms exist, and that the ones we have documented are exceptional not in their ages but in their detectability (e.g., due to epistasis among sites). Moreover, because we restrict our attention to biallelic SNPs, we will have missed structural and multiallelic variation, which may be more common targets of balancing selection than point mutations. Nonetheless, the convergence of independent lines of evidence supports a clear conclusion. A TSP between humans and chimpanzees, long treated as a gold standard for balancing selection, is by itself insufficient evidence, and in most cases may simply be the footprint of neutral, incomplete lineage sorting.

## Acknowledgments

We thank Felix Wu for organizing the sequencing of the chimpanzee samples; we are also grateful to Trevor Cousins, Omer Gokumen, William Milligan, Jeff Wall as well as Anastasia Stolyarova and other members of the Przeworski lab for helpful discussions.

## Data Availability

Sequencing datasets used are documented in Supplementary Table 3: *Newly sequenced chimpanzee pedigree samples* and Supplementary Table 4: *Summary of all chimpanzee samples used*.

Posterior probabilities of coalescence by time points >2 Mya, from an earlier version of Cousins et al. and provided to us via personal correspondence.

Code to reproduce all analyses and figures are available at https://github.com/hmunby/human-chimp-tsp/

## Methods

### Chimpanzee samples

We used publicly available whole-genome sequences for chimpanzees, to which we added data obtained by the Przeworski lab in 2020 from blood samples for a pedigree of ten chimpanzees (including five founders), which were provided in 2005 when MP was employed at Brown University by the Primate Foundation of Arizona. DNA libraries were prepared for sequencing by Genewiz using a PCR-free protocol and sequenced on Illumina HiSeq 2500 with 2 x 150 cycles to mean fold-coverage of between 16 and 44x in 2020 (Supplementary Table 3). Sequencing data for these samples are available on NCBI’s SRA under the project accession PRJNA1357362.

### Chimpanzee datasets and variant calling

We downloaded SRA files of chimpanzee whole genome sequences from Venn et al. 2014 [51], Tatsumoto et. al. 2017 [52], Besenbacher et al. 2019 [53] and de Manuel et al. 2016 [50] and combined them with the 10 whole genome sequences that we had generated. In total, there are 59 samples, of which 43 are unrelated (i.e., not first- and second-degree relatives) (Supplementary Table 4). For each sample, reads were aligned to the PanTro6 assembly with bwa-mem in paired-end mode. Duplicates were marked in the resulting BAM files with GATK’s MarkDuplicates. We then called variants with GATK (version 4.2.3) following GATK best practices. Specifically, for each chromosome, we generated individual gVCFs with HaplotypeCaller before consolidating all samples with GenomicsDBImport and jointly calling variants with GenotypeGVCFs. After removing indels and multi-allelic sites, we were left with 43,481,990 SNPs.

### Filtering

Following GATK 4 best practices, we retained sites with Phred Quality Score (QUAL) > 30, QualByDepth (QD) > 2, RMSMappingQuality (MQ) > 40, MappingQualityRankSumTest (MQRankSum) < 6, FisherStrand bias (FS) < 60 and ReadPosRankSum < 6. We removed any sites where the total read depth across samples is <0.5-fold or >1.5-fold the median of the whole-genome average coverage per site. We excluded sites with excess heterozygosity by retaining only sites with ExcessHet < 28.69 (i.e. with p-values corresponding to more than 3 standard deviations away from Hardy-Weinberg equilibrium). Finally, we removed any sites where ≥10% of samples are missing genotypes. After this filtering, we were left with 38,169,920 SNPs.

We used the pedigrees in our full dataset to identify sites with Mendelian errors using PLINK (--mendel). Any sites exhibiting one or more Mendelian violations in our total sample were removed (a further 862,705 SNPs). After this analysis, we retained only founders from the pedigrees and wild-born individuals from de Manuel et al. (2016) for downstream analyses.

The reference assembly to which we mapped reads may contain regions where paralogs have been collapsed into a single copy. Differences between such copies could potentially generate false heterozygote calls, affecting our identification of shared variants. We therefore ran BISER [74] on both the PanTro6 and the telomere-to-telomere (T2T) assembly in order to detect segmental duplications that are collapsed in PanTro6 but are resolved in the T2T assembly. A total of 9707 regions, comprising 11.3% of the genome, were identified as duplicated by this approach. We removed the 4,878,666 SNPs (13.1%) that fall in these cryptic segmental duplications, thereby retaining 32,428,549 SNPs.

### Sub-species identification

We retrieved the subspecies labels of all publicly available samples from their respective publications (Supplementary Table 4). In order to identify the subspecies of the founder individuals in our newly sequenced pedigree, we performed principal component analysis (PCA) on the total set of 59 chimpanzee samples. We used PLINK to LD-prune the variant set, i.e., to reduce linkage disequilibrium levels among SNPs (--indep-pairwise 50 10 0.1), and perform the PCA. We found that the first two principal components separate all unrelated samples into three clusters, corresponding to subspecies labels *Pan troglodytes verus* (Western), *Pan troglodytes schweinfurthii* (Eastern) and *Pan troglodytes troglodytes* (Central). Of the five founder individuals in our newly sequenced pedigree, four clustered with the labelled Western samples and the other one clustered with the Eastern samples (see Supplementary Figure 1). Our final set of 43 unrelated samples thus consists of 14 Eastern, 14 Central and 15 Western chimpanzees.

### Liftover of chimpanzee variants

To map the positions onto the human genome, the filtered VCFs were lifted to hg19 using bcftools +liftover plugin [75]. We removed sites without a unique mapping in the new reference. Sites where neither the reference or alternative allele called in chimpanzee match the hg19 reference became multi-allelic and therefore were also removed. As a further check, we lifted the remaining variants back to the PanTro6 reference and removed any SNPs that did not map back to their original position. Lastly, we required that the region +/-100bp around each chimpanzee SNP maps also uniquely to hg19.

### Human (1000GP) data processing

We downloaded the 1000 Genomes Project dataset of short variant calls in the Phase 3 dataset 2504 individuals. We removed indels and multiallelic sites to generate a set of biallelic SNPs. To remove SNPs in potential paralogs, as we had done for the chimpanzee data, we used two published datasets: the set of segmental duplications identified using the human T2T genome [76] and copy-number variants called in 1000GP Phase 3 [46]. Any SNPs falling within these regions were removed for the shared variant analysis (i.e., 16.0% of the initial set).

### Identification of shared variants between humans and chimpanzees

We considered only human SNPs with a minor allele frequency ≥5% in 1000GP and chimpanzee SNPs with a minor allele count of three or more in our unrelated sample. To identify shared variants, we intersected the human and chimpanzee VCFs using *bcftools isec*. We then removed any sites that are multi-allelic in the resulting output; such sites represent positions where both species harbour a biallelic polymorphism but the segregating alleles differ (e.g. A/T in human vs. A/G in chimpanzee). The resulting final output is a set of 66,650 SNPs at which the same two alleles are segregating in human and chimpanzee samples. We used the Genome Reference Consortium’s definition of the MHC locus (chr6:28,477,797-33,448,354) to identify MHC variants and excluded them from downstream analyses where noted.

### Allele age estimates

To characterize the ages of shared variants, we used allele-age estimates from two local genealogical reconstruction methods applied to the 1000 Genomes Project (1000GP) Phase 3 data, *Relate* [47] and *SINGER* [48].

*Relate* infers the topologies of local genealogies along the genome from the full 1000GP dataset and estimates branch lengths separately for each of the 26 subpopulations, yielding population-specific allele-age estimates [47]. We used estimates for the Yoruba in Ibadan, Nigeria (YRI, n = 108), chosen because it is among the largest 1000GP samples, is one of the more genetically diverse populations, and because Leffler et al. relied on a YRI sample, aiding comparison. *Relate* YRI estimates were available for 56,468 shared SNPs (85%).

*SINGER* reconstructs the ancestral recombination graph (ARG) and samples from its posterior, allowing the uncertainty in each local genealogy to be quantified. Deng et al. applied *SINGER* to a sample of 200 African haplotypes from 1000GP and generated 100 posterior samples of the ARG [48]. Age estimates were available for 19,009 shared SNPs (29%). Although *SINGER* provides estimates for far fewer of our SNPs, it appears less prone than *Relate* to the downward bias affecting age estimates for the oldest alleles [54] and reports a posterior distribution of genealogies rather than a single fixed topology; we therefore show *SINGER*-based results alongside *Relate* in the Supplement (Supplementary Figure 2).

For each 1000GP variant, both methods provide the ages of the start and end of the genealogical branch on which the derived allele is inferred to have arisen (for *SINGER*, across all 100 posterior ARG samples). To proceed, we assumed that the mutation is uniformly distributed along its branch. Where a single point estimate is required, for example to define the candidate set for genealogical reconstruction (see below) or to bin SNPs by age (Fig 1B), we take the branch midpoint as a point estimate. To compare age distributions between SNP sets (Fig 1A), we instead computed, for a given set, the expected number of SNPs older than an age a as the sum over SNPs of the probability that each is older than *a* (under the uniform-branch assumption, the fraction of its branch that lies above *a*), evaluated at regular age intervals. We applied this approach to the shared SNPs and to all common 1000GP SNPs (MAF ≥ 5%), converting generations to years by assuming a generation time of 29 years.

#### Bias of the branch-midpoint age estimator

We estimated the age of an allele by its branch midpoint. To characterise the bias in this point estimator, we simulated neutral genealogies under a constant size model using msprime [77]. We simulated 1 Mb segments without recombination under a constant-size neutral coalescent (msprime; 2,000 diploid samples, i.e. 4,000 haplotypes; effective population size Nₑ = 10,000; mutation rate μ = 1×10⁻⁸ per site per generation). For every mutation, we recorded its true age together with the midpoint age of the branch on which it arose, defined as the mean of the true start and end dates of the branch, excluding mutations on the root branch and those at a derived-allele frequency below 1%. Simulation replicates were run until approximately 10⁸ mutations had been recorded (∼5×10⁴ replicates).

### Local genealogical reconstruction around old shared variants

To identify trans-species polymorphisms, we focused on the shared SNPs old enough to plausibly have persisted IBD since the human–chimpanzee common ancestor. We retained as candidates all shared SNPs whose estimated age as provided by *Relate* exceeds 4 Mya, excluding the MHC region, namely 860 SNPs. We set this threshold below the inferred human–chimpanzee split time (∼5–6 Mya) deliberately, as age estimates for the oldest variants are biased downward (see Results and Supplementary Figure 3).

For each candidate, we reconstructed the local genealogy of a combined sample of humans and chimpanzees. We first re-called chimpanzee variants in the region surrounding each candidate, defined by grouping candidate SNPs within 100 kb of one another and extending the interval by 250 kb on either side. Chimpanzee reads for all samples were mapped to the hg19 reference and variants called with GATK as described above, applying the same filtering criteria used for the PanTro6 calls. The resulting chimpanzee genotypes were phased with SHAPEIT2 and restricted to unrelated individuals (n = 43). For each region, the chimpanzee VCF was merged with the corresponding 1000GP VCF using bcftools merge, and any site that became multiallelic (i.e. was segregating for different alleles in the two species) was removed.

Because *SINGER* scales to samples on the order of hundreds of haplotypes, and in order to obtain a combined sample with comparable numbers of individuals per species, we downsampled the human genotypes to 50 individuals per region. To avoid distorting the genealogy at the focal SNP, we performed this downsampling so as to preserve its allele frequency, sampling individuals from each diploid genotype class at the focal SNP in the same proportions as in the full 1000GP dataset. We then ran *SINGER* on each merged region with an effective population size (Ne) of 50,000 and a mutation rate per bp per generation of μ = 1.25 × 10⁻⁸ [78,79], for 10,000 MCMC iterations; the first 4,000 were discarded as burn-in and we then retained every 60th iteration to yield 100 posterior samples of the ARG. Because thinning does not fully remove autocorrelation between samples drawn from a single MCMC chain, we repeated this procedure three times per genomic region using independent random seeds, giving 300 posterior samples in total. ARG samples were converted to tree-sequence format using *SINGER*’s convert_to_tskit, and all downstream operations and visualization used tskit [80,81].

Following the expectations for trans-species balanced polymorphisms in Gao et al. [36], we assessed support for each candidate from the local tree topology at its position (for SNP positions with multiple mutations, we used the oldest; Supplementary Figure 4). We required, first, that when the focal mutation arose, the genealogy contained more than one but no more than three lineages (i.e. the mutation is not on the root branch), and second, that the first coalescent event following the mutation joined a branch whose descendants are all chimpanzee haplotypes to a branch whose descendants are all human haplotypes. For each candidate, we recorded the proportion of the 300 posterior ARG samples meeting both criteria as a measure of support for it being a trans-species polymorphism (TSP). We classified a candidate as a TSP if it met both topology criteria in more than 80% of the 300 posterior ARG samples. This approach yielded a final set of 117 TSPs, whose positions were lifted to GRCh38 (liftOver) for downstream annotation. Because the ARG-reconstruction sample combines haplotypes from two species, the choice of Ne is not straight-forward. We therefore repeated the reconstruction and TSP classification for Ne=20,000 and Ne=100,000, using a single chain in each case, which yielded 111 and 133 TSPs respectively. All of the qualitative results that are reported in the text are insensitive to this choice.

Of the 860 candidates, we were able to place the candidate SNP on a reconstructed local genealogy and evaluate it as a TSP in 802 cases (93.3%). The remaining 58 SNPs were lost at the same step, the chimpanzee remapping and recalling on hg19. These SNPs were called in the original chimpanzee variant calls (made against the chimpanzee reference and lifted to hg19) but not when chimpanzee reads were re-mapped directly onto the human reference; at sites where the chimpanzee sequence diverges from human or mappability is low, reads either fail to align to hg19 or align with low mapping quality, leaving too few confidently-placed reads for the caller to emit a genotype. Of these 58 SNPs, 27 were not near any SNP that was successfully remapped. The remaining 31 were within clusters of candidate SNPs (i.e., <100kb from another candidate); thus, while we were unable to test these specific SNPs, we were able to reconstruct their local genealogies and test for TSP at nearby SNPs.

When analysing SNPs by region (Table 1), we grouped clusters of TSPs falling within 10 kb of one another. The 117 TSPs formed 59 regions, 44 singletons and 15 regions containing more than one TSP (Table 1). To confirm that the TSPs grouped into a region correspond to a single segregating haplotype in humans, rather than independently evolving sets of variants, we computed pairwise genotypic linkage disequilibrium (r²; PLINK) among the TSPs in the 1000 Genomes Project phase 3 reference panel, computing r² separately within the African (AFR, n = 661) and European (EUR, n = 503) samples. In almost every region with more than one TSPs, they are in strong pairwise LD (minimum r² ≥ 0.8 in both samples), effectively forming a single LD block. In the IGFBP7 region the TSPs resolve into two separate LD blocks, containing 2 and 4 TSPs respectively, a finding that has been previously reported [36,38]. In the LINC01676 region the TSPs form a single block in AFR (minimum r² = 0.81) but are less tightly linked in EUR (0.58).

### Measures of background selection

We characterised the local strength of linked (background) selection using a map of the background-selection coefficient B for autosomes (the CADD best-fit B-map [64]), where low B indicates strong background selection. The map was lifted to GRCh38 and mean values for each 1 kb window of the genome were calculated. For analyses where we stratify the genome by background selection, we binned the genome by quantiles of B in 1 kb windows across autosomes (with decile 1 / quartile 1 corresponding to the strongest background selection). The same 1 kb window edges were used throughout: to assign each TSP region its BMAP decile, to construct matched sets of loci for enrichment analyses, and when looking at regions that remain uncoalesced in the human-chimpanzee ancestor (Figures 4 and 6).

### Functional annotations of shared and trans-species variants

#### Variant Effects

We annotated SNPs with their variant effects using snpEff. Because snpEff reports one effect per overlapping transcript, each SNP was assigned the single, most severe consequence under a fixed consequence ranking; protein-truncating effects (stop-gained, stop-lost, start-lost) were collapsed to "LOF". Coding and regulatory categories were ordered by the estimates of functional constraint in Agarwal et al. ([82]; Fig 2B), and the non-coding categories that they did not consider (intergenic, intron, intragenic, non-coding-transcript exon) were considered to have the least consequence. In the plot (Figure 3), 5′ and 3′ UTR variants were combined ("UTR") as were upstream/downstream variants (within 5 kb of a gene) ("flanking"). Percentages for each category for each SNP set are reported with binomial standard errors.

#### Sequence conservation

PhyloP conservation scores were extracted at each SNP by exact coordinate (hg19). As a primary measure, we used scores from Fu et al. 2014 [61], computed on the Ensembl Compara 36-eutherian-mammal EPO_LOW_COVERAGE alignment with the human sequence excluded (see main text). As a comparison, we also used the UCSC phyloP 46-way placental-mammal track and recovered qualitatively similar results. Empirical cumulative distributions were compared across sets; differences between the set of TSPs and other SNP sets were tested with the Kolmogorov–Smirnov test.

#### Overlap with candidate cis-regulatory elements

SNP positions (GRCh38) were intersected (bedtools) with the ENCODE SCREEN registry of candidate cis-regulatory elements (GRCh38, SCREEN V4; 2,348,854 cCREs [83]). We reported the fraction of each SNP set overlapping any cCRE.

#### Gene assignment and functional enrichment of trans-species loci

Each region was assigned a single gene using a tiered rule that aggregates evidence across all of its TSPs, applied through the Open Targets Genetics GraphQL API [84,85]. In order of priority: (i) a GWAS locus-to-gene (L2G) effector-gene prediction with score ≥ 0.5; otherwise (ii) a gene supported by a colocalising molecular QTL (eQTL, sQTL or pQTL) credible set, choosing among multiple molQTL genes by the number of tissues, then modalities, then credible sets; and otherwise (iii) the nearest protein-coding gene (GENCODE v49, bedtools closest). The 59 regions and their assigned genes are given in Table 1 (for regions with more than one TSP) and Supplementary Table 2 (all regions). Because several regions are assigned the same gene, these 59 regions correspond to 55 distinct genes, and all counts below are of distinct genes.

We then tested for functional enrichments among these genes using Gene Ontology (Biological Process, Cellular Component, Molecular Function 2023) [86], KEGG (2021) [87], Reactome (2022) [88], and MSigDB Hallmark (2020) [89] categories, obtained from the Enrichr gene-set library collection [90,91], as well as UniProt-keyword KW-0325 glycoproteins and membrane glycoproteins (the intersection of genes which are annotated as UniProt-keyword KW-0325 glycoprotein and UniProt-keyword KW-0472 membrane) [92]. For every category containing at least two of the TSP genes, we performed two different enrichment tests. First, within each category, we tabulated the number of genes to which TSPs were assigned (“TSP genes”). We compared this overlap to a background of all autosomal protein-coding genes using a one-sided hypergeometric test, thereby asking whether more of the TSP genes fall in the category than expected when the same number of genes is drawn at random from all autosomal protein-coding genes. Because this approach does not account for the genomic properties of the TSP genes, we next tested enrichment of these same categories in the TSP genes against 2,000 matched gene sets (described below). Specifically, we compared the proportion of TSP genes falling in the category to the corresponding proportion in each matched set, and computed an empirical p-value of (1 + b) / (1 + 2000), where b is the number of the matched gene sets whose proportion was at least as large as observed in the TSP genes. P-values were corrected by the Benjamini–Hochberg procedure across categories [93].

To generate the 2,000 matched sets of genes, we considered the strength of background selection as well as the number of TSPs in a region. Specifically, for each TSP region, the matching procedure was as follows. We calculated the mean BMAP value of SNPs in the TSP region and then identified all common autosomal SNPs (1000GP, MAF ≥ 5%) with a BMAP value within 20 of that mean; we then sampled 2,000 of these SNPs without replacement. We built a matched region around each of these 2,000 SNPs by taking the *n* nearest common SNPs, where *n* is the total number of SNPs in the TSP region minus one. We recorded the genomic spans of the resulting matched regions, which were comparable to, but slightly smaller than, the TSP regions. Each of these matched regions was then assigned to a gene through a pipeline identical to that used to assign the TSP regions to genes. With this procedure, we generated the 2,000 matched sets, each comprising the distinct genes assigned to 59 matched regions. Given that gene-set libraries are highly redundant, to aid in interpretation of our enrichment results, we grouped significant categories by the overlap of their contributing genes, joining categories whose combined coefficient (0.5 x Jaccard + 0.5 x overlap coefficient) reached 0.375 (default in EnrichmentMap; [95]).

### Coalescent inference and decoding

To identify regions of the genome that remain uncoalesced around the time of the human-chimpanzee split, we applied the approach of Cousins et al. [45] to both species. For humans, we used the estimates of Cousins et al. directly: the authors applied PSMC, as implemented in *cobraa* (github.com/trevorcousins/cobraa), to seven individuals of recent African ancestry (one from each African subpopulation of 1000GP), and used the posterior decoding (step size 1 kb) to obtain, for every 1 kb segment and individual, the posterior probability of coalescence within each PSMC time interval [60]. The posterior probability that a segment has not coalesced by a given time boundary is 1 minus the summed probability of coalescence in all more recent intervals. The authors kindly provided us with the estimated probabilities of remaining uncoalesced for every time boundary older than 2 Mya for every 1 kb segment.

To replicate this analysis in chimpanzees, we selected nine samples, three from each subspecies. For each sample, VCFs were generated from the PanTro6-aligned BAMs (bcftools mpileup -q 20 -Q 20 -C 50, then bamCaller,py from *msmc-tools* (github.com/stschiff/msmc-tools)) and converted to the PSMC input multihetsep (MHS) file format with generate_multihetsep.py from *msmc-tools.* Low-mappability sites were filtered out using a PanTro6 mappability mask generated with SNPable (k-mer 150 bp, stringency 75%). We ran PSMC, as implemented in *cobraa*, with D = 64, b = 100, spread1 = 0.075, spread2 = 100 and a mutation-to-recombination ratio of 1.5, fixing the scaled mutation rate at θ = 0.001 across all samples. Using the inferred PSMC parameters and the input MHS files, we then ran posterior decoding for each individual using *cobraa*’s decode mode with a step size of 1 kb, obtaining coalescence posterior probabilities within each time interval for each segment. To obtain the posterior probability of a segment having coalesced by a specific timepoint, we summed the probabilities of coalescence across all previous time intervals (i.e., between that time point and the present). The posterior probability of an individual’s two haplotypes remaining uncoalesced beyond that timepoint is given by 1 minus the probability of having coalesced. Time steps are in coalescent units, so to convert to years, we assumed a generation time of 24 years [94] and mutation rate of 1.27×10⁻⁸ per bp per generation [53]. We focus on 5.5 Mya, which is around the peak of the coincident increase in effective population sizes in humans and chimpanzees inferred by Cousins et al. [45]. We take the first timepoint to exceed this value, which falls at 5,755,625 years in chimpanzees; the corresponding value in humans is 5,589,889 years.

To pinpoint the regions that remain uncoalesced >5.5 Mya, we summed the posterior probabilities of remaining uncoalesced across samples within a species for each 1 kb region. We then retained segments for which the sum exceeds 1, thus identifying segments where in expectation more than one sampled individual carries an uncoalesced haplotype pair. To do so in chimpanzees, we first lifted segments from PanTro6 to the human reference.

### Regions uncoalesced in humans and chimpanzees

We considered the intersection of the segments that were uncoalesced by 5.5 Mya in humans and in chimpanzees. For each background selection decile, we then calculated the proportion of sequence callable in both species that is also uncoalesced in both. The analysis was conducted for all autosomes, but with the MHC excluded. For each of the four SNP sets, 1000GP SNPs with a MAF ≥ 5%, all shared SNPs, old shared (≥4 Mya), and TSPs, we computed the fraction of SNPs that occurs in regions uncoalesced by 5.5. Mya in both species with bedtools intersect (the results are shown in Figure 2).

#### Testing whether regions of deep coalescences in humans and chimpanzees co-occur more than expected under neutrality and purifying selection alone

For each BMAP quartile, we computed the length of the callable genome that is uncoalesced by 5.5 Mya. To focus on unconstrained segments, we considered the subset of sites with a mean phyloP (UCSC 100-way vertebrate track) below 0.332, corresponding to the 90th percentile of the autosomal distribution of mean phyloP scores across 1 kb windows. We tried cutoffs of 0.090 (70th percentile), 0.150 (80th percentile) and 0.617 (95th percentile) and the qualitative conclusions were unchanged. We also excluded the MHC region from this analysis. 95% confidence intervals were obtained by bootstrap resampling the intersected segments with replacement (200 replicates) and taking the 2.5th and 97.5th percentiles of the fraction across replicates.

#### Gene enrichment in regions uncoalesced in both species

We merged flanking segments that are uncoalesced in both species into contiguous regions. We then calculated the mean BMAP value of each of these contiguous regions across sites. Regions that are physically close may still not be entirely independent; for example, a longer segment could have been fragmented by liftOver between assemblies, or because the intervening regions narrowly miss our criterion to be called as uncoalesced in one of the species. Therefore, in addition to only merging regions if separated by 0 kb, we also tried merging regions separated by distances of 5 kb, 10 kb or 25 kb.

To test for gene enrichments among the uncoalesced regions, we assigned each one to its closest protein-coding gene (GENCODE v49; autosomes only) using bedtools closest. Where more than one gene was equidistant, one was chosen at random, and because that choice is arbitrary, the analysis was repeated over 200 independent random resolutions. Regions were stratified by BMAP quartile, resulting in a list of genes for each quartile that we tested for enrichment in two ways.

(1) We compared the genes in the test set to a background of genes in the same quartile. To obtain the set of background genes, we mapped every segment of the genome jointly callable in humans and chimpanzees in that BMAP quartile to its closest protein-coding gene and took the resulting unique set of genes. Then, in each BMAP quartile and for each gene category, we took the number of genes annotated to that category in the uncoalesced regions (k_test_) and in the background (k_bg_), as well as the total number of genes for each (n_test_ and n_bg_) and calculated the fold enrichment as (k_test_ / n_test_) / (k_bg_ / n_bg_). We assessed the significance of observing at least k_test_ genes when n_test_ are drawn from the n_bg_ background genes with a one-sided hypergeometric test. We excluded all gene assignments within the MHC locus in both test and background sets.
(2) We tested for enrichment accounting for gene length and the distance to the nearest other gene. This second approach aims to overcome a possible confounding effect of enrichment tests, namely that when each segment is assigned to its closest gene, large genes (considering their total length, including introns) or genes farther from other genes are assigned a larger portion of the genome, and so have more segments assigned to them by chance. If genes with these features are enriched for a specific functional category, this assignment procedure may lead to a spurious enrichment for that category. Mindful of this issue, we tested against resampled null gene sets as follows. For each observed uncoalesced region, we randomly drew a contiguous block of the jointly-callable genome spanning the same number of bps from within the same BMAP quartile, and assigned it to its closest gene. By this approach, large or more isolated genes are drawn more often. Repeating this procedure 10,000 times yields a null distribution for the proportion of genes in each gene category, matched for genomic features. We reported the enrichment of a category as the proportion of observed genes in the category divided by the mean proportion across null sets, with a permutation p-value of (1+b) / (1 + 10,000), where b is the number of null sets whose proportion was at least as large as observed. As in the hypergeometric test, we tested only categories with at least three observed genes and applied the Benjamini-Hochberg correction across categories within each quartile. Gene assignments within the MHC were again excluded. We followed the same approach to cluster categories with overlapping genes as for categories enriched among TSP genes. As a further sensitivity analysis, we also repeated the entire procedure discarding assignments where the closest gene lay further than 50 kb or 100 kb, applying the same cutoff to null sets; qualitative results were unchanged.

By these approaches, we identified membrane glycoprotein as enriched the strongest background selection quartile (see Results) regardless of the choice of distance over which to merge segments or the exclusion of regions further than 50 kb of a protein-coding gene. Other gene categories were less stable. For example, some olfactory receptor categories were enriched but lost genes for larger merge distances, while the anti-fungal immunity category no longer survived the multiple-testing correction.

## Supplementary Figures & Tables

**Supplementary Table 1:** Per-SNP annotations for the 117 TSPs. One row per TSP, ordered by chromosome and position. *Position and alleles*: Coordinates in the human genome are given for the hg19/GRCh37 assembly and also lifted to GRCh38. We provide rsIDs for all variants for which one is available. Reference and alternative alleles are with respect to hg19. Chimpanzee coordinates and alleles are with respect to panTro6 (GCA_002880755.3). *Sequence context*: CpG status; background-selection value (BMAP, scaled 0–1000, higher values indicating weaker background selection); Roulette mutation rate; and phyloP score from the 36-eutherian-mammal EPO alignment with the human sequence masked (see Methods, "Functional annotation of shared and trans-species variants"). *Predicted effect*: snpEff annotation and VEP most severe consequence. *Gene assignment*: nearest protein-coding gene and distance to it; the assigned gene and the tier of evidence supporting the assignment (locus-to-gene, molecular QTL, or nearest); whether the variant is present in the Open Targets index; the number of credible sets; molecular QTL types and tissues; and GWAS locus-to-gene effector genes with their scores in parentheses (Methods, "Gene assignment and functional enrichment of trans-species loci"). *Allele age:* Lower and upper bounds and midbranch age of the polymorphism in the YRI population as inferred by *Relate* [47]. *TSP support*: The percentage of the 300 posterior genealogies, pooled across three independent chains, in which the SNP met our trans-species topology criteria (see Methods, “Identification of trans-species polymorphisms”).

| Region (hg19) | Gene | Assignment | TSPs | Physical span (bp) | Genetic span (x10<sup>-3</sup> cM) | Variant types | BMAP decile | Literature | GWAS (L2G effector) |
| --- | --- | --- | --- | --- | --- | --- | --- | --- | --- |
| chr21:41,205,924-41,208,874 | HMGN1 | molqtl | 12 | 2950 | 0.72 | intergenic | 9 | Bitarello et al. 2018 | — |
| chr5:8,017,470-8,024,003 | MTRR | molqtl | 11 | 6533 | 3.251 | intergenic | 9 | Leffler et al. 2013; Bitarello et al. 2018 | — |
| chr4:144,655,489-144,662,054 | GYPB | l2g | 8 | 6565 | 2.243 | intron | 4 | Bitarello et al. 2018; Leffler et al. 2013; Rocha et al. 2026 | Lactate levels; Total bilirubin levels (+20 more) |
| chr4:115,378,668-115,379,737 | UGT8 | l2g | 7 | 1069 | 0.021 | intergenic | 7 | Bitarello et al. 2018 | Calcium levels; Urolithiasis |
| chr4:57,918,296-57,919,705 | IGFBP7 | l2g | 6 | 196 / 484 | 0.565 / 1.245* | intron | 9 | Leffler et al. 2013; Bitarello et al. 2018; DeGiorgio et al. 2014; Rocha et al. 2026 | Height |
| chr4:71,210,148-71,210,970 | MUC7 | molqtl | 5 | 822 | 0.222 | intergenic | 7 | Bitarello et al. 2018 | — |
| chr2:1,138,967-1,139,647 | SH3YL1<sup>†</sup> | molqtl | 4 | 680 | 1.434 | intron | 8 | Bitarello et al. 2018; DeGiorgio et al. 2014; Rocha et al. 2026 | — |
| chr1:106,125,135-106,131,147 | LINC01676 | molqtl | 3 | 6012 | 3.899 | downstream gene, intergenic | 8 | — | Body mass index; Weight (+1 more) |
| chr11:24,791,431-24,792,057 | LUZP2 | nearest | 3 | 626 | 0.009 | intron | 10 | Bitarello et al. 2018; DeGiorgio et al. 2014 | — |
| chr2:237,968,214-237,968,421 | COPS8 | nearest | 3 | 207 | 0.058 | upstream gene | 10 | — | — |
| chr4:56,144,663-56,147,967 | SRD5A3 | molqtl | 3 | 3304 | 0.667 | intergenic | 4 | Leffler et al. 2013 | — |
| chr5:16,919,819-16,919,992 | MYO10 | l2g | 2 | 173 | 0.133 | upstream gene | 10 | — | Heel bone mineral density |
| chr10:25,593,349-25,593,541 | GPR158 | nearest | 2 | 192 | 0.101 | intron | 8 | Rocha et al. 2026 | — |
| chr10:67,278,595-67,279,509 | CTNNA3 | l2g | 2 | 914 | 0.001 | intergenic | 7 | Bitarello et al. 2018 | Testosterone levels; Total testosterone levels (+8 more) |
| chr5:29,637,400-29,637,695 | CDH6 | nearest | 2 | 295 | 0.221 | intergenic | 9 | — | — |
| chr2:224,638,996-224,638,996 | AP1S3 | l2g | 1 | 0 | 0 | intron | 9 | — | Arm predicted mass left; Basal metabolic rate |
| chr1:246,380,688-246,380,688 | SMYD3 | nearest | 1 | 0 | 0 | intron | 5 | — | — |
| chr4:12,087,957-12,087,957 | HS3ST1 | nearest | 1 | 0 | 0 | intergenic | 8 | — | — |
| chr2:224,887,004-224,887,004 | SERPINE2 | nearest | 1 | 0 | 0 | intron | 8 | — | — |
| chr4:57,698,600-57,698,600 | SPINK2 | nearest | 1 | 0 | 0 | intergenic | 8 | — | — |
| chr2:129,359,232-129,359,232 | HS6ST1 | nearest | 1 | 0 | 0 | intergenic | 10 | — | — |
| chr2:42,237,228-42,237,228 | PKDCC | l2g | 1 | 0 | 0 | intergenic | 3 | — | Ankle spacing width; Femoral neck bone mineral density (+9 more) |
| chr2:35,952,400-35,952,400 | CRIM1 | nearest | 1 | 0 | 0 | upstream gene | 7 | — | — |
| chr1:247,627,001-247,627,001 | AHCTF1 | molqtl | 1 | 0 | 0 | intergenic | 7 | Bitarello et al. 2018 | Mean platelet (thrombocyte) volume |
| chr1:116,732,423-116,732,423 | MAB21L3 | nearest | 1 | 0 | 0 | intergenic | 8 | Bitarello et al. 2018 | — |
| chr1:5,348,639-5,348,639 | AJAP1 | nearest | 1 | 0 | 0 | intergenic | 10 | — | — |
| chr1:4,917,460-4,917,460 | AJAP1 | nearest | 1 | 0 | 0 | intergenic | 10 | — | — |
| chr4:162,038,353-162,038,353 | FSTL5 | nearest | 1 | 0 | 0 | intergenic | 7 | — | — |
| chr4:141,921,518-141,921,518 | RNF150 | molqtl | 1 | 0 | 0 | upstream gene | 8 | Rasmussen et al. 2014; DeGiorgio et al. 2014 | — |
| chr4:58,570,867-58,570,867 | IGFBP7 | nearest | 1 | 0 | 0 | intergenic | 3 | Bitarello et al. 2018 | — |
| chr4:71,386,175-71,386,175 | MUC7 | l2g | 1 | 0 | 0 | intron | 6 | — | Uterine fibroids |
| chr4:58,495,926-58,495,926 | IGFBP7 | nearest | 1 | 0 | 0 | intron | 3 | — | — |
| chr3:145,043,291-145,043,291 | PLOD2 | nearest | 1 | 0 | 0 | intergenic | 8 | — | — |
| chr4:32,646,855-32,646,855 | PCDH7 | nearest | 1 | 0 | 0 | intergenic | 7 | — | — |
| chr4:180,069,715-180,069,715 | AGA | nearest | 1 | 0 | 0 | intergenic | 9 | Bitarello et al. 2018 | — |
| chr7:36,547,245-36,547,245 | AOAH | nearest | 1 | 0 | 0 | intergenic | 10 | — | — |
| chr8:21,455,324-21,455,324 | GFRA2 | l2g | 1 | 0 | 0 | intergenic | 8 | — | Body mass index |
| chr8:74,978,724-74,978,724 | LY96 | nearest | 1 | 0 | 0 | intron | 7 | — | — |
| chr8:54,528,520-54,528,520 | ATP6V1H | nearest | 1 | 0 | 0 | intergenic | 6 | — | HDL cholesterol levels x long total sleep time interaction |
| chr9:7,215,204-7,215,204 | KDM4C | molqtl | 1 | 0 | 0 | intergenic | 7 | — | — |
| chr4:172,136,342-172,136,342 | GALNTL6 | nearest | 1 | 0 | 0 | intergenic | 8 | — | — |
| chr6:166,673,913-166,673,913 | MPC1 | molqtl | 1 | 0 | 0 | intergenic | 10 | — | — |
| chr4:165,753,488-165,753,488 | MARCHF1 | molqtl | 1 | 0 | 0 | intergenic | 10 | — | — |
| chr10:62,623,360-62,623,360 | RHOBTB1 | molqtl | 1 | 0 | 0 | intergenic | 7 | — | — |
| chr10:59,743,375-59,743,375 | IPMK | nearest | 1 | 0 | 0 | downstream gene | 7 | Bitarello et al. 2018 | — |
| chr10:26,184,344-26,184,344 | MYO3A | nearest | 1 | 0 | 0 | intergenic | 4 | Bitarello et al. 2018 | — |
| chr10:132,883,127-132,883,127 | TCERG1L | nearest | 1 | 0 | 0 | intergenic | 10 | — | — |
| chr11:133,338,872-133,338,872 | OPCML | l2g | 1 | 0 | 0 | intron | 6 | — | Substance use disorder |
| chr13:37,973,335-37,973,335 | POSTN | nearest | 1 | 0 | 0 | intergenic | 9 | Bitarello et al. 2018 | — |
| chr13:59,433,547-59,433,547 | DIAPH3 | nearest | 1 | 0 | 0 | intergenic | 3 | — | Body mass index; Body fat percentage (+9 more) |
| chr11:5,373,738-5,373,738 | OR51E2 | molqtl | 1 | 0 | 0 | upstream gene | 8 | — | — |
| chr13:74,300,726-74,300,726 | KLF12 | l2g | 1 | 0 | 0 | intron | 6 | — | Adolescent idiopathic scoliosis; Standing height |
| chr14:22,758,552-22,758,552 | OXA1L | molqtl | 1 | 0 | 0 | intergenic | 10 | — | — |
| chr15:54,984,861-54,984,861 | UNC13C | nearest | 1 | 0 | 0 | intergenic | 8 | — | — |
| chr14:89,538,401-89,538,401 | FOXN3 | nearest | 1 | 0 | 0 | intergenic | 8 | — | — |
| chr16:5,605,114-5,605,114 | RBFOX1 | l2g | 1 | 0 | 0 | intron | 10 | DeGiorgio et al. 2014 | Weight, inverse-rank normalized |
| chr17:58,829,124-58,829,124 | BCAS3 | l2g | 1 | 0 | 0 | upstream gene | 1 | — | Creatinine levels; Glutamine levels (+7 more) |
| chr17:70,908,606-70,908,606 | SLC39A11 | nearest | 1 | 0 | 0 | intron | 10 | — | — |
| chr20:52,817,810-52,817,810 | PFDN4 | l2g | 1 | 0 | 0 | intergenic | 10 | — | Colorectal cancer (oestrogen-progestogen hormone therapy interaction) |

**Supplementary Table 2:** Regions with TSPs. Summary of the 59 regions in which the 117 TSPs occur. The column labeled “region” gives the genomic coordinates (hg19) of each cluster of TSPs, with SNPs within 10 kb assigned to the same region. “Gene” refers to the gene that the region is assigned to based on supporting evidence tiers (see Methods) and “Assignment” details which of these tiers we use for the assignment. “TSP count” gives the number of shared SNPs in this region that are estimated to be ≥4Mya and which meet our TSP topology criterion in >80% of ARG samples. “Physical span” gives the maximum distance between genomic coordinates of TSPs in the region, and “Genetic span” provides the genetic distance, based on HapMap II [95]. “Annotation” refers to the SnpEff consequence class of the TSPs within the region. “BMAP decile” refers to the decile of the background selection map in which the region lies (1 = strongest, 10 = weakest background selection). “Literature” indicates prior studies that report the region as a long-term balancing-selection or TSP candidate. “GWAS (L2G effector)” lists genome-wide association study (GWAS) traits for which a TSP in the region is the index variant of a GWAS credible set and the assigned gene is the Open Targets locus-to-gene effector. Regions with multiple TSPs are also described in Table 1. **\***The IGFBP7 region is composed of two linkage groups (with 2 and 4 SNPs respectively); we report the spans of each of these linkage groups separately. **†** We assign the TSPs in this region to the gene *SH3YL1* based on molecular QTL evidence but note that the TSPs lie within the gene *SNTG2* which is reported by DeGiorgio et al. (2014) and Rocha et al. (2026) [28,58].

| Sample ID | Mean fold-coverage (after mapping) | Sex | Pedigree founder | Subspecies |
| --- | --- | --- | --- | --- |
| 1031 | 43.68 | Female | Yes | *P. t. verus* (Western) |
| 1034 | 40.67 | Female | Yes | *P. t. verus* (Western) |
| 1059 | 27.37 | Female | Yes | *P. t. verus* (Western) |
| 1064 | 28.74 | Female | Yes | *P. t. verus* (Western) |
| 1076 | 17.53 | Female | No | Hybrid backcross<br>Western x F1 |
| 1078 | 15.67 | Female | No | Hybrid backcross<br>Western x F1 |
| 2003 | 38.85 | Male | Yes | *P. t. schweinfurthii*<br>(Eastern) |
| 2047 | 28.05 | Male | No | F1 hybrid -<br>Western x Eastern |
| 2059 | 27.28 | Male | No | F1 hybrid<br>Western x Eastern |
| 2073 | 29.76 | Male | No | F1 hybrid<br>Western x Eastern |

**Supplementary Table 3.** Newly sequenced chimpanzee pedigree samples. The column “Sample ID” gives the sample identifier; “Mean fold-coverage” gives the mean number of reads mapped to a site in the PanTro6 reference; “Sex” gives the sex of the individual; “Pedigree founder” indicates where the individual was one of the founders of the pedigree (Yes) or if it is offspring of other individuals in the pedigree (No); “Subspecies” refers to which of the four subspecies of chimpanzee the individual belongs to and states which individuals are hybrids based on their parentage.

| Citation | NCBI Project ID | Pedigree | Samples used | Subspecies | Total N | Unrelated N |
| --- | --- | --- | --- | --- | --- | --- |
| de Manuel *et al.* (2016) | PR JEB15086 | No | SAMEA4374773, SAMEA4374799, SAMEA4374766, SAMEA4374797, SAMEA4374778, SAMEA4374775, SAMEA4374780, SAMEA4374782, SAMEA4374772, SAMEA4374763, SAMEA4374781, SAMEA4374787, SAMEA4374777, SAMEA4374770, SAMEA4374767, SAMEA4374791, SAMEA4374765, SAMEA4374769, SAMEA4374776, SAMEA4374785, SAMEA4374788, SAMEA4374764, SAMEA4374771, SAMEA4374790, SAMEA4374792, SAMEA4374774, SAMEA4374794, SAMEA4374796, SAMEA4374795, SAMEA4374798, SAMEA4374793, SAMEA4374768, SAMEA4374779, SAMEA4374789 | Western, Eastern, Central | 34 | 32 |
| Venn *et al.* (2014) | PRJEB5937 | Yes | SAMEA2421540, SAMEA2421541, SAMEA2421542, SAMEA2421543, SAMEA2421544, SAMEA2421545, | Western | 9 | 3 |
|  |  |  | SAMEA2421546, SAMEA2421547,<br>SAMEA2421548 |  |  |  |
| Tatsumoto *et al.* (2017) | PRJDB3537 | Yes | SAMD00026329, SAMD00026330,<br>SAMD00026331 | Western | 3 | 2 |
| Besenbacher *et al.* (2019) | PRJEB29710 | Yes | SAMEA5204226, SAMEA5204228,<br>SAMEA5204229 | Western | 3 | 1 |
| This study | PRJNA1357362 | Yes | 1031, 1034, 1059, 1064, 1076,<br>1078, 2003, 2047, 2059, 2073 | Western,<br>Eastern, Hybrid | 10 | 5 |
| Total | - | - | - |  | 59 | 43 |

**Supplementary Table 4. Summary of all chimpanzee samples used.**

**Supplementary Figure 1:**
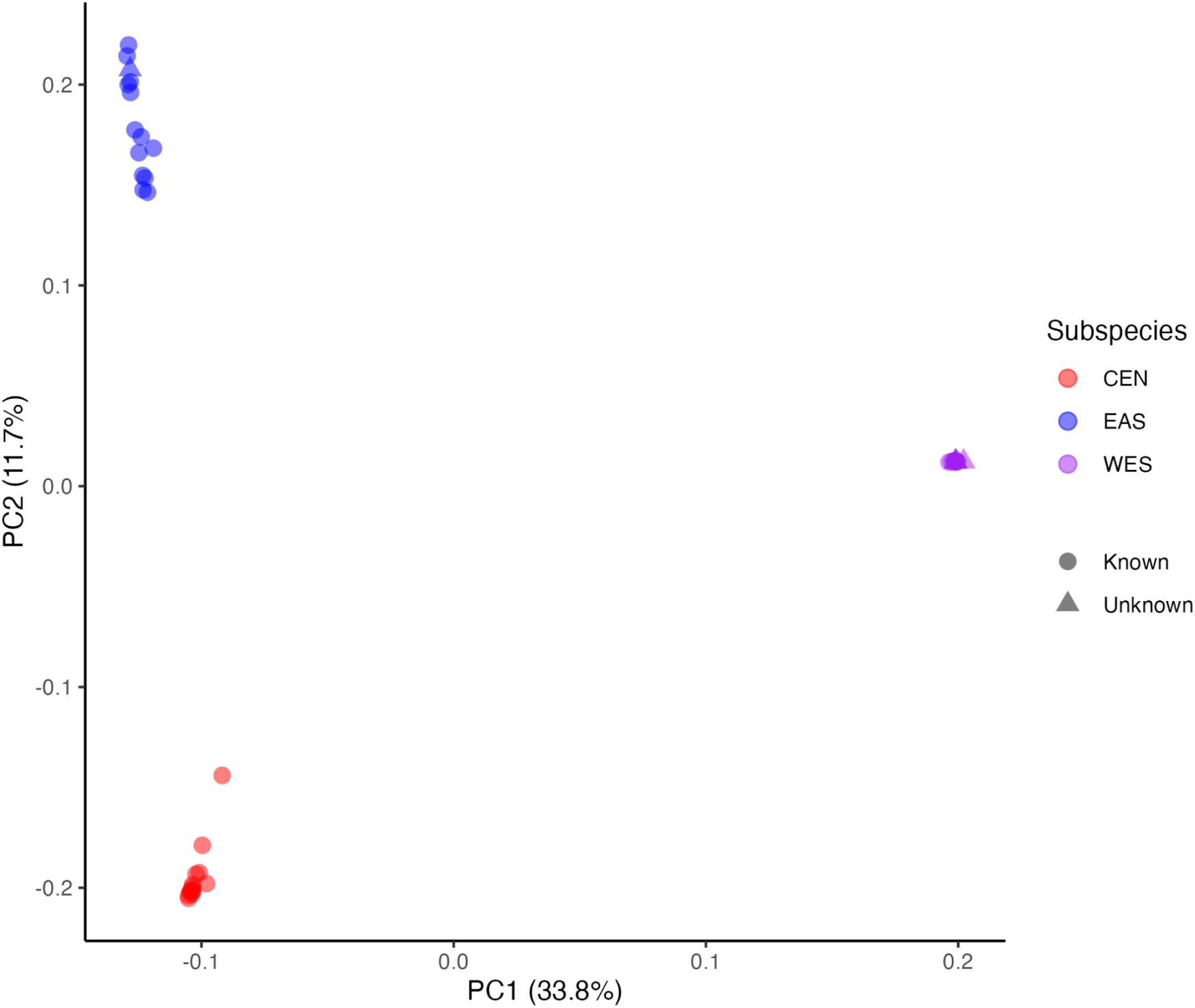
First two Principal Components from the Principal Component Analysis (PCA) of the unrelated chimpanzee samples. The PCA is based on filtered and LD-pruned, whole-genome SNP data for 43 unrelated chimpanzees (see Methods). Shown are the primary axis of variation (PC1) and secondary axis of variation (PC2), with the proportion of variance explained in parenthesis. Samples are colored according to their subspecies labels (Central = red, Eastern = blue, Western = purple). Shapes indicate whether the subspecies label was previously assigned (circles) or generated in this study (triangles). As can be seen, four of the five samples for which we collected genome sequence data cluster with samples previously labelled as Western and one with Eastern samples.

**Supplementary Figure 2:**
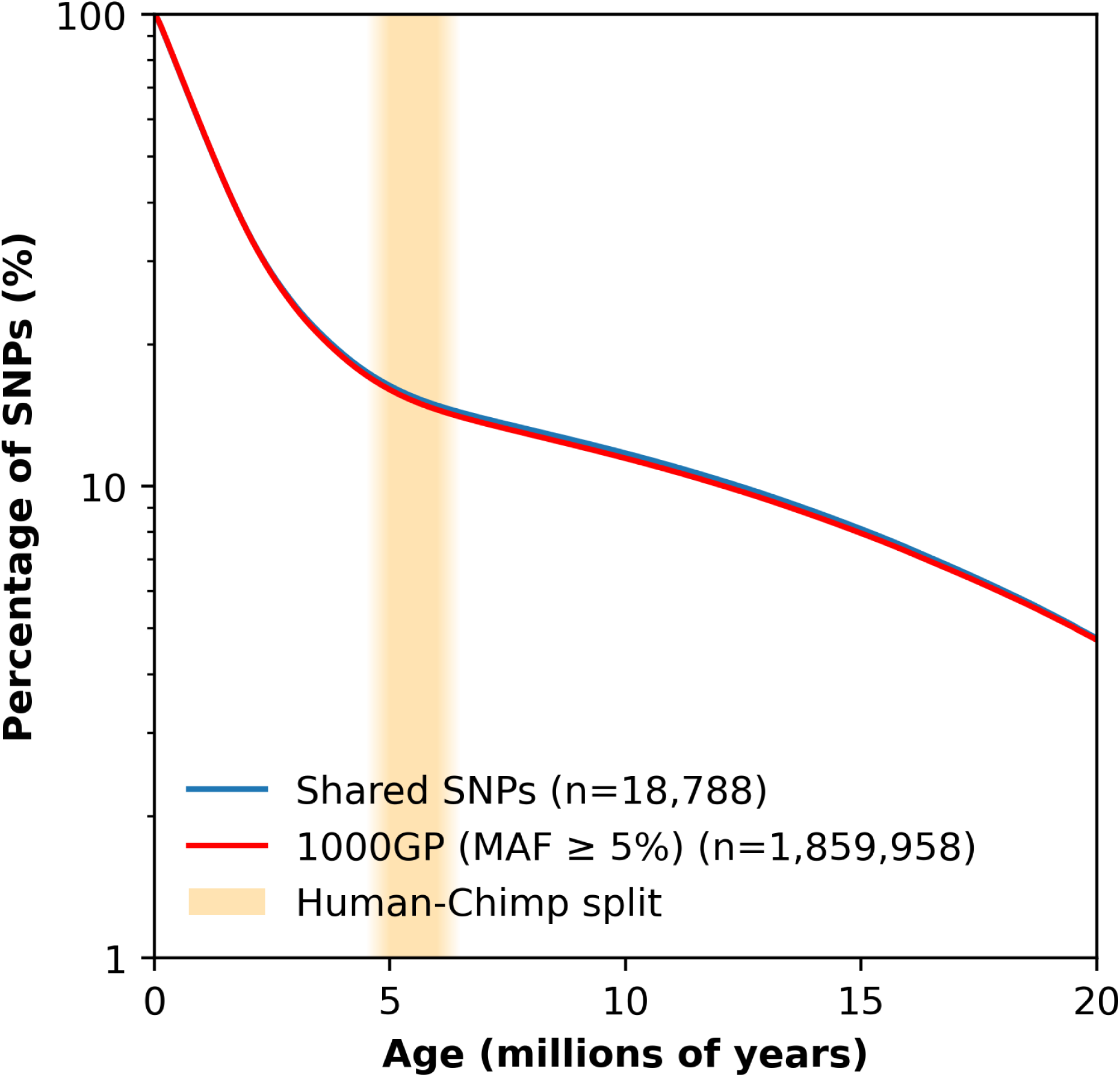
Allele ages of shared SNPs and common human variants, estimated by *SINGER*. In blue are estimated ages of SNPs shared between humans and chimpanzees (n = 18,788) and and in red that of genome-wide common variants (1000 Genomes Project, MAF ≥ 5%; n = 1,859,958), plotted as the percentage of SNPs estimated to be older than a given age (on a log scale). The shaded band marks the human and chimpanzee species are thought to have split off from one another (∼5–6 Mya). As expected from previous work, *SINGER* infers a much higher proportion of SNPs in both sets to be very old, as compared to *Relate* [47,48].

**Supplementary Figure 3:**
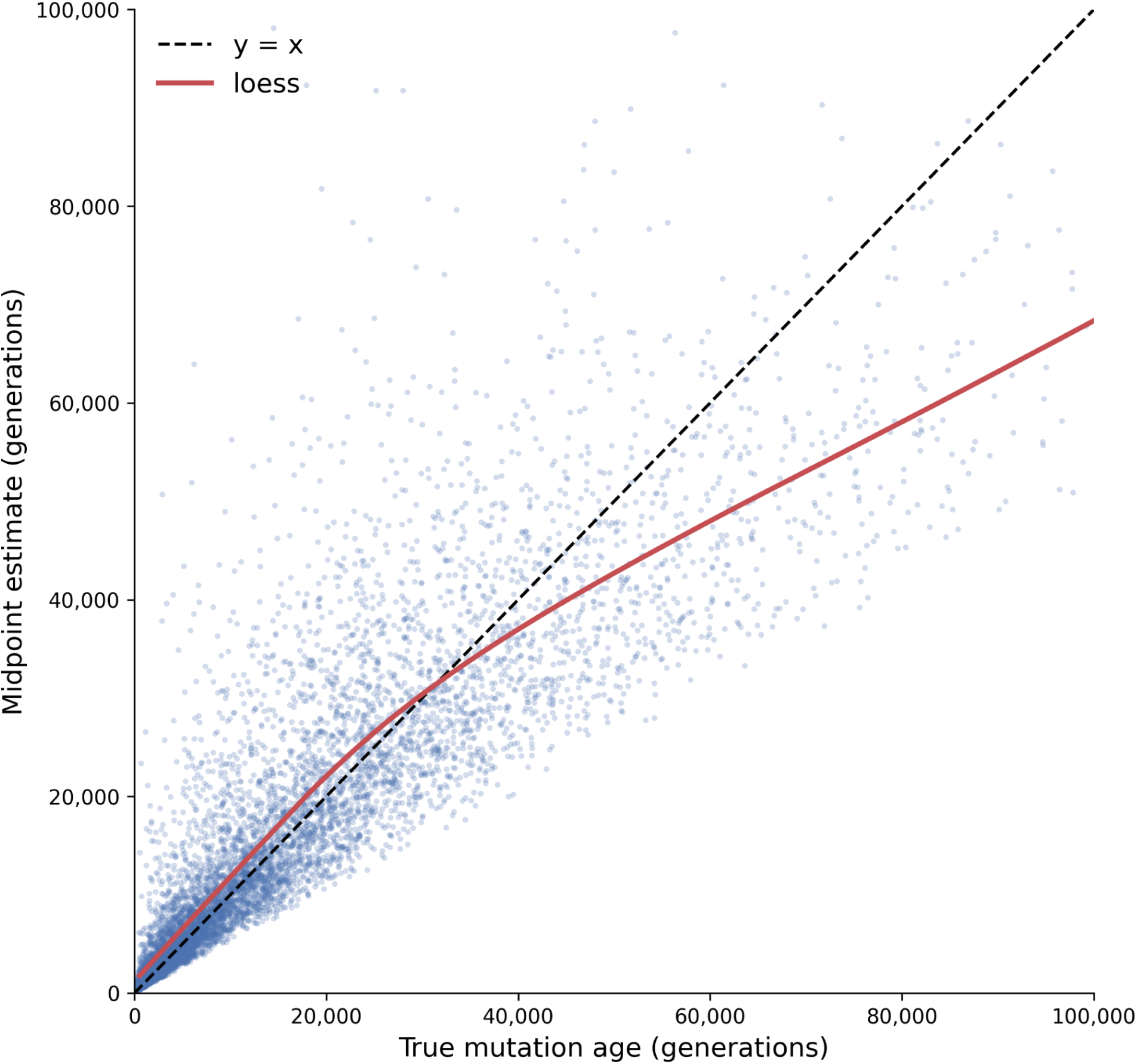
Bias in the mid-branch age estimator of mutation age, estimated from neutral coalescent simulations. We simulated neutral genealogies under a coalescent model using msprime, for 2000 diploid samples, a constant effective population size Ne = 10,000, and mutation rate μ = 10⁻⁸ per site per generation. We ran approximately 50,000 simulations to yield 100,090,793 mutations on non-root branches with a derived-allele frequency ≥ 1%; for each mutation we recorded the true age together with the midpoint age of its branch. Points show the midpoint estimate against the true age for a random subsample of 10,000 mutations, in generations. The dashed line is what would be expected for an unbiased estimator (y = x) and the red curve is a loess of the midpoint estimate on the true age, computed over all simulated mutations. As can be seen, in this demographic setting, this estimator is downwards biased for mutations older than ∼3-4Ne generations.

**Supplementary Figure 4:**
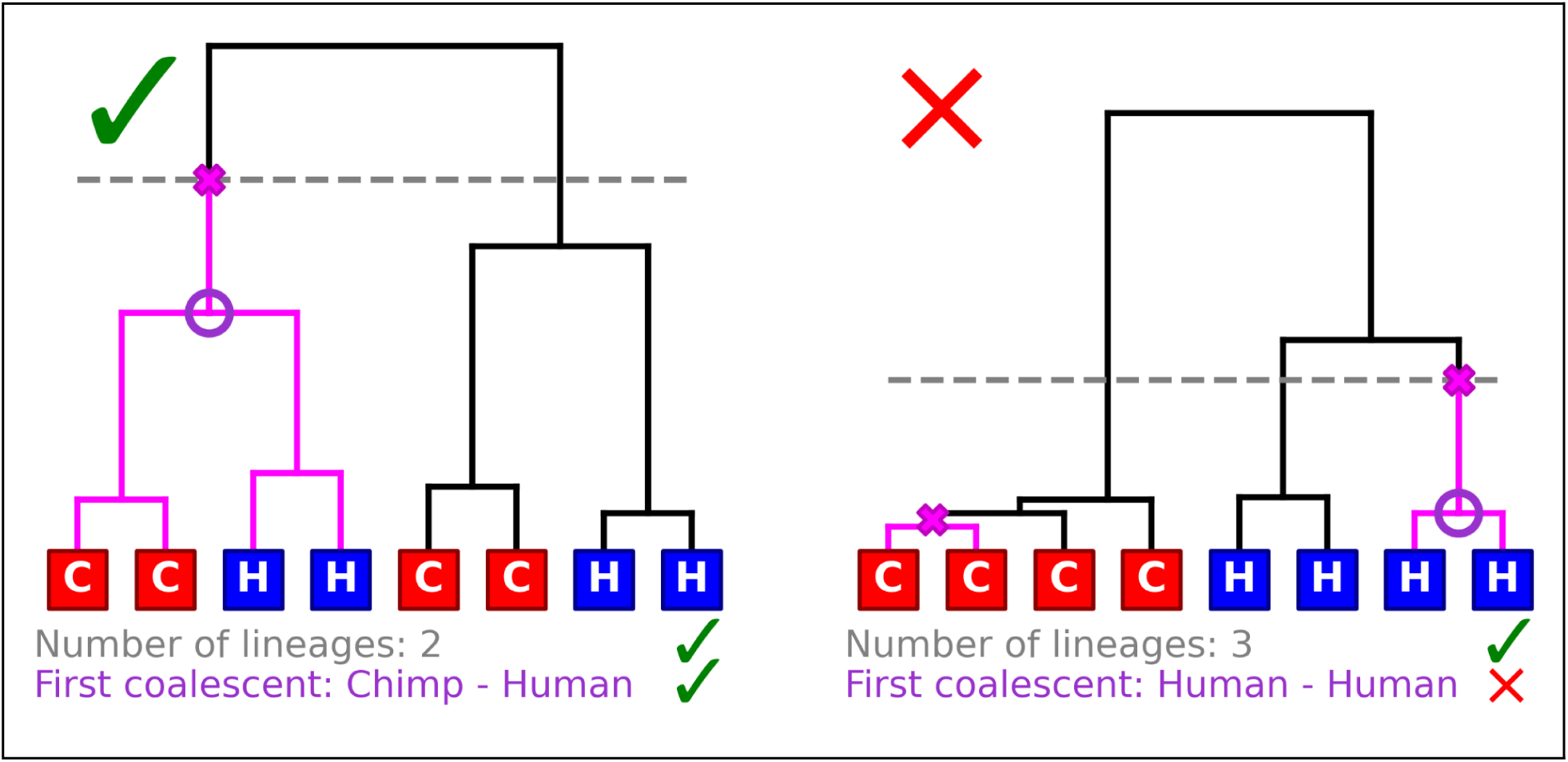
Criteria applied to local genealogies to evaluate the evidence for a TSP. The example of a reconstructed genealogy of eight haplotypes illustrates the criteria that we use to evaluate if a shared variant is likely a TSP. We require lineages from different species (blue vs. red tips) that both carry the derived (or ancestral) allele (pink branches) to coalesce with one another before they coalesce with lineages of their own species carrying the ancestral (derived) allele. Specifically, we require that, at the time of the oldest mutation at the focal site, there are either two or three lineages segregating in the genealogy (grey dashed line). We further consider whether the first coalescent below this mutation is between a lineage whose descendants are all humans and a lineage whose descendants are all chimpanzees (purple circle). On the left is an example of a polymorphism that meets both of these criteria. On the right, we show a possible reconstruction for a mutation that arose independently in the two species; the mutation in humans is very old and meets our first criterion but, because all of its descendants are human haplotypes, it does not meet the second one.

**Supplementary Figure 5:**
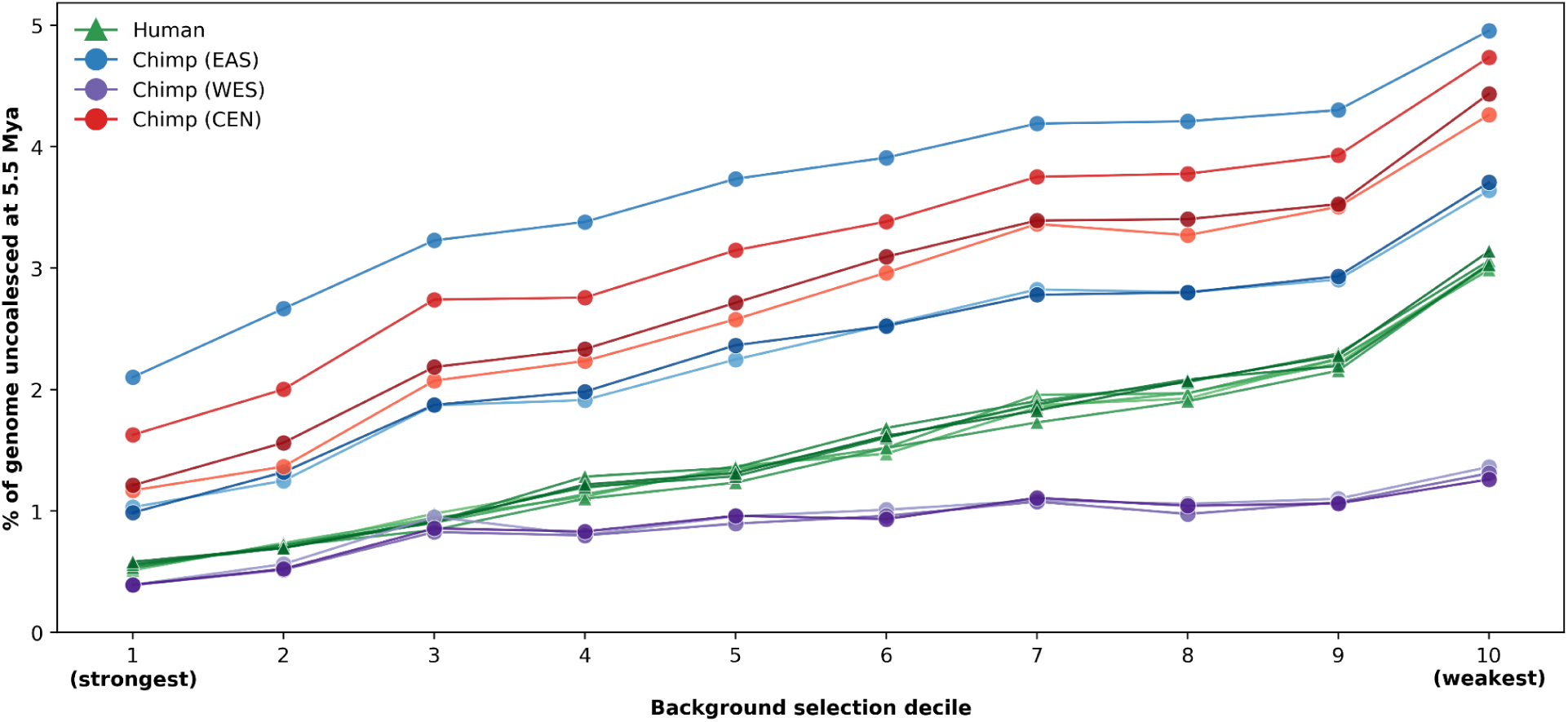
Expected percentage of 1 kb segments of the genome that remain uncoalesced by 5.5 Mya, stratified by the strength of background selection, in chimpanzees and humans. The autosomal genome was partitioned into deciles of BMAP, a measure of the strength of background selection (see Methods). The MHC region is excluded. On the y-axis is the expected number of 1kb segments within that decile that are inferred to be uncoalesced 5.5 Mya in each individual (see Methods). Results are shown for the posterior decoding of PSMC run on seven humans (green triangles) (taken from [96]) and nine chimpanzees (circles), three from each of three subspecies: Eastern in blue, Western in purple, and Central in red. As expected, populations/subspecies with greater diversity show a higher fraction of the genome uncoalesced by 5.5 Mya.

**Supplementary Figure 6:**
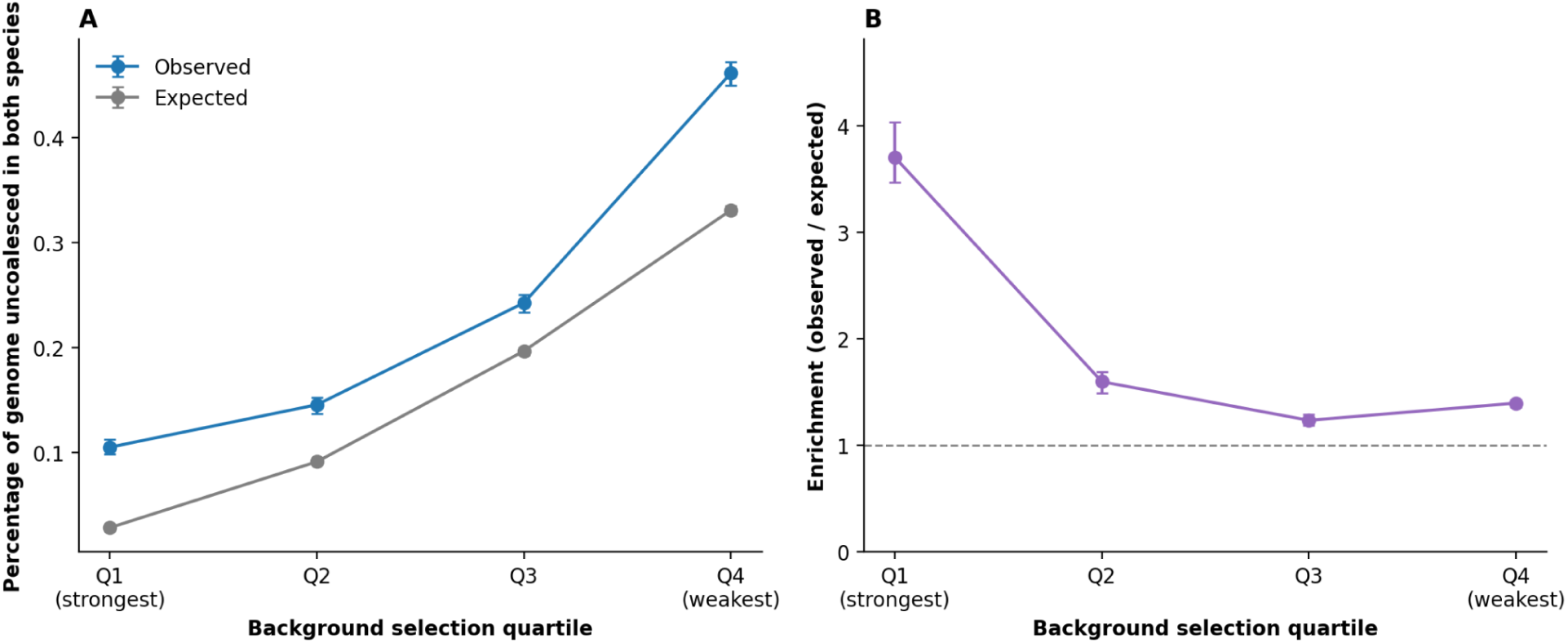
Regions uncoalesced in both species relative to what would be expected if independent, stratified by the strength of background selection, including the MHC. Autosomal regions were split into quartiles, based on the strength of background selection. Constrained segments were excluded. (**A**) For each quartile, we report the estimated proportion of the sequence that is uncoalesced in both humans and chimpanzees (in blue) by 5.5 Mya versus what we would expect assuming that coalescent events are independent in the two species (see Methods; in grey). (**B**) The ratio of observed to expected proportions, as a measure of enrichment. Points are estimates and error bars 95% bootstrap confidence intervals.

